# Retrotransposon-mediated variation of a chitin synthase gene confers insect resistance to *Bacillus thuringiensis* Vip3Aa toxin

**DOI:** 10.1101/2024.02.18.580648

**Authors:** Zhenxing Liu, Chongyu Liao, Luming Zou, Minghui Jin, Yinxue Shan, Yudong Quan, Hui Yao, Lei Zhang, Peng Wang, Zhuangzhuang Liu, Na Wang, Anjing Li, Kaiyu Liu, David G. Heckel, Kongming Wu, Yutao Xiao

## Abstract

*Bacillus thuringiensis* (*Bt*) crops expressing Vip3Aa are highly efficacious in controlling major lepidopteran pests and delaying evolution of pest resistance. Although practical resistance to Vip3Aa in the field has not been reported, to proactively manage the pest resistance, there is an urgent need to better understand the genetic basis of resistance to Vip3Aa. This is particularly important for the fall armyworm (*Spodoptera frugiperda*), one of the most destructive pests around the world, which has evolved practical resistance to *Bt* crystal (Cry) toxins. Here, a highly Vip3Aa-resistant (resistance ratio: 5,562-fold) strain of *S. frugiperda* was selected in the laboratory. Results from bulked segregant analysis, fine-scale mapping, and genetic linkage analysis indicate that a mutation in the midgut-specific chitin synthase gene, *SfCHS2*, is strongly associated with high-level resistance to Vip3Aa. The resistance is ascribed to the transcriptional variation caused by retrotransposon insertion. The same variation of *SfCHS2* was also detected in a field population. Importantly, knockout of *SfCHS2* via CRISPR/Cas9 in susceptible *S. frugiperda* confers its complete resistance (>10,000-fold) to Vip3Aa. Also, we demonstrate that knockout of *CHS2* can result in complete resistance to Vip3Aa in additional lepidopteran species, suggesting a general role of this gene in Vip3Aa resistance among lepidopteran pests. These results reported here would contribute to monitor and management of pest resistance to Vip3Aa.

## Introduction

Insecticidal proteins from the bacterium *Bacillus thuringiensis* (*Bt*) have protected transgenic crops covering more than a billion hectares from pest damage since 1996 (1). In addition to protection from feeding damage, benefits include suppression of pest populations and consequent reduced damage to non-transgenic crops (2, 3), reduced use of chemical insecticides and compatibility with integrated pest management including biological control by natural enemies (4, 5). However, as Bt cotton became more widely used in countries with different agricultural practices, and as Cry toxins were deployed in additional crops such as maize and soybean, Cry toxin resistance has begun to evolve in situations where not all the assumptions of the original resistance management plan were met (6). More than 26 practical cases of resistance that decrease the efficacy of Bt crops have been documented in some populations of 11 pest species across seven countries (6).

To counter resistance to Cry toxins, farmers have increased the proportion of pyramid crops that produce Bt vegetative insecticidal protein Vip3Aa and Cry proteins simultaneously (7-9). Vip3Aa insecticidal proteins are expressed in the vegetative stage of growth starting at mid-log phase as well as during sporulation and showed toxicity against multiple lepidopteran insect species including black cutworm (*Agrotis ipsilon*), fall armyworm (*Spodoptera frugiperda*), beet armyworm (*Spodoptera exigua*), tobacco budworm (*Heliothis virescens*), and corn earworm (*Helicoverpa zea*) (10). Vip3Aa was considered as a novel insecticidal protein produced by Bt, since it can facilitate pore formation in larvae midgut via a mode of action distinct with that of Cry proteins (11-14). The Vip3Aa-transgenic crops have been approved for commercial cultivation around for more than 10 years (15). Practical resistance to Vip3Aa in the field has not been reported previously. However, since 2012, field or laboratory-evolved resistance to Vip3A toxin in various target pests has been reported successively around the world (12, 13, 16-21). Especially, early warnings of field-evolved resistance to Vip3Aa have been reported for two major pests (6, 8, 22). Better understanding of mechanism of resistance to Bt proteins has been confirmed to be helpful for resistance management in the field (23). However, knowledge about the genetic basis of resistance to Vip3Aa toxin in lepidopteran pests is still largely unknown. To proactively manage the pest resistance, it is urgently needed to have more knowledge about the genetic basis of resistance to Vip3Aa.

The knowledge about genetic basis of resistance to Vip3Aa is especially important for the fall armyworm (*S. frugiperda*). As an invasive pest, *S. frugiperda* has recently expanded its traditional range in North and South America to the rest of the world except continental Europe and Antarctica (24). Its high fecundity and migratory tendencies pose serious problems for agriculture in the newly colonized regions (25). It has evolved practical resistance to Bt crops producing Cry toxins (6). Previous study have identified multiple potential receptors for Vip3Aa or proteins involved in tolerance of pest against Vip3Aa toxicity (26-29). However, up to now, there is no evidence to link those proteins with resistance to Vip3Aa selected in the laboratory or field. Although a number of Vip3Aa-resistant insect colonies have been established and the genetic of Vip3Aa resistance have been well characterized (12, 13, 17, 18, 20, 30), the genetic basis associated with Vip3Aa resistance for those cases have not been reported.

Recently, downregulation of a transcription factor, *SfMyb*, was reported to be associated with resistance to Vip3Aa in *S. frugiperda* (31). However, the variation of *SfMyb* can causes just 206-fold resistance to Vip3Aa in *S. frugiperda*. Yet the genetic basis involved in high-levels resistance to Vip3Aa, like as those resistance cases reported previously (12, 20, 32), has not been reported as far as we know. Here, we report the identification of association, for the first time, between high-level resistance to Vip3Aa in *S. frugiperda* and alternative splicing of a midgut-specific chitin synthase caused by retrotransposon insertion. This resistance allele with retrotransposon insertion was also detected in the field. Furthermore, we demonstrate that knockout of this gene via CRISPR/Cas9 can confer complete resistance of *S. frugiperda* to Vip3Aa toxin. The effect was also discovered in other two related insect pests. Our results strongly support that this midgut-specific chitin synthase plays a substantial role in the mode of action of Vip3Aa toxin.

## Results

### Characterization of a highly Vip3Aa-resistant strain of *S. frugiperda*

A highly Vip3Aa-resistant strain (Sfru_R3) of *S. frugiperda* was established in the laboratory from a susceptible strain (SS), collected from a maize field in China (Ruili, Yunnan Province) in late 2019, after 17 generations of selection using diet incorporated with gradually increased concentration of Vip3Aa protoxin (Table S1). Results from bioassay with artificial diet showed that Sfru_R3 exhibited a more than 5562-fold resistance ratio relative to the SS (Table 1). In the case of bioassay with transgenic plants, the survival rate of larvae from Sfru_R3 feeding with Vip3Aa-transgenic maize was 23.33%, while no survivor was observed for the larvae from SS (Table S2). Bioassay of progeny generated from the reciprocal crosses between Sfru_R3 and SS showed that the high-level Vip3Aa resistance was an autosomal and recessive (*D* = -0.78, *D*_*ML*_ = 0.065) trait (Table S3). Besides, no cross-resistance to Cry1Ab, Cry1Ac, Cry1F and Cry2Ab was detected in Sfru_R3 (Table S4). Susceptibility tests of progeny from back-crosses (*χ*^*2*^ = 3.52, *P* > 0.05) and F2 generation (*χ*^*2*^ = 1.68, *P* >0.05) suggest that Vip3Aa resistance in Sfru_R3 could be controlled by single genetic locus (Table S5).

**Table 1.** Bioassay results for Vip3Aa-susceptible, resistant strains.

| Insect strain | N<sup>a</sup> | EC<sub>50</sub> (95% CI) (µg/cm<sup>2</sup>)<sup>b</sup> | Slope ± SE | χ<sup>2</sup> | df | Resistance ratio <sup>c</sup> |
| --- | --- | --- | --- | --- | --- | --- |
| SS | 144 | 0.077 (0.062 - 0.092) | 4.72 ± 0.91 | 0.59 | 3 | - |
| Sfru_R3 | 64 | 428.26 (312.41 - 616.95) | 4.37 ± 1.12 | 0.25 | 2 | 5562 |
| F<sub>1</sub>: Sfru_R3♀<br>× SS♂ | 144 | 0.27 (0.23-0.32) | 5.94 ± 1.11 | 1.52 | 3 | 3.5 |
| F<sub>1</sub>: Sfru_R3♂<br>× SS♀ | 144 | 0.26 (0.14 - 0.55) | 3.75 ± 0.55 | 6.75 | 3 | 3.4 |
| pooled F<sub>2</sub> | 2304 | 0.33 (0.20 - 0.94) | 1.11 ± 0.12 | 131.9 | 3 | 4.3 |
| pooled BC to<br>SS | 4608 | 2.97 (1.04 - 9.52) | 1.40 ± 0.082 | 5.15 | 5 | 38.6 |

<sup>a</sup> Total number of neonates tested.
<sup>b</sup> Median effective concentration (EC<sub>50</sub>) causing 50% of neonates (first-instar) to die or fail to advance to the third instar in the 7-day test period.
<sup>c</sup> Resistance ratio for a population equals its EC<sub>50</sub> divided by the EC<sub>50</sub> of SS.

### Identification of candidate gene genetically linked with resistance to Vip3Aa toxin

To identify the genetic basis associated with Vip3Aa resistance in Sfru_R3, a bulked segregant analysis was performed using genomic DNA from the parental moths, Vip3Aa-resistant (F2-R) and Vip3Aa-susceptible (F2-S) larvae from F2 progeny (Fig. S1). A genomic region ranging from 6.47 to 10.30 Mb in chromosome 1 (Chr. 1) was mapped, due to the ΔSNP index for SNPs in this region being all over the threshold value (Figure 1A and B). Subsequently, the ∼4 Mb genomic region was selected for further fine-scale mapping.

**Figure 1.**
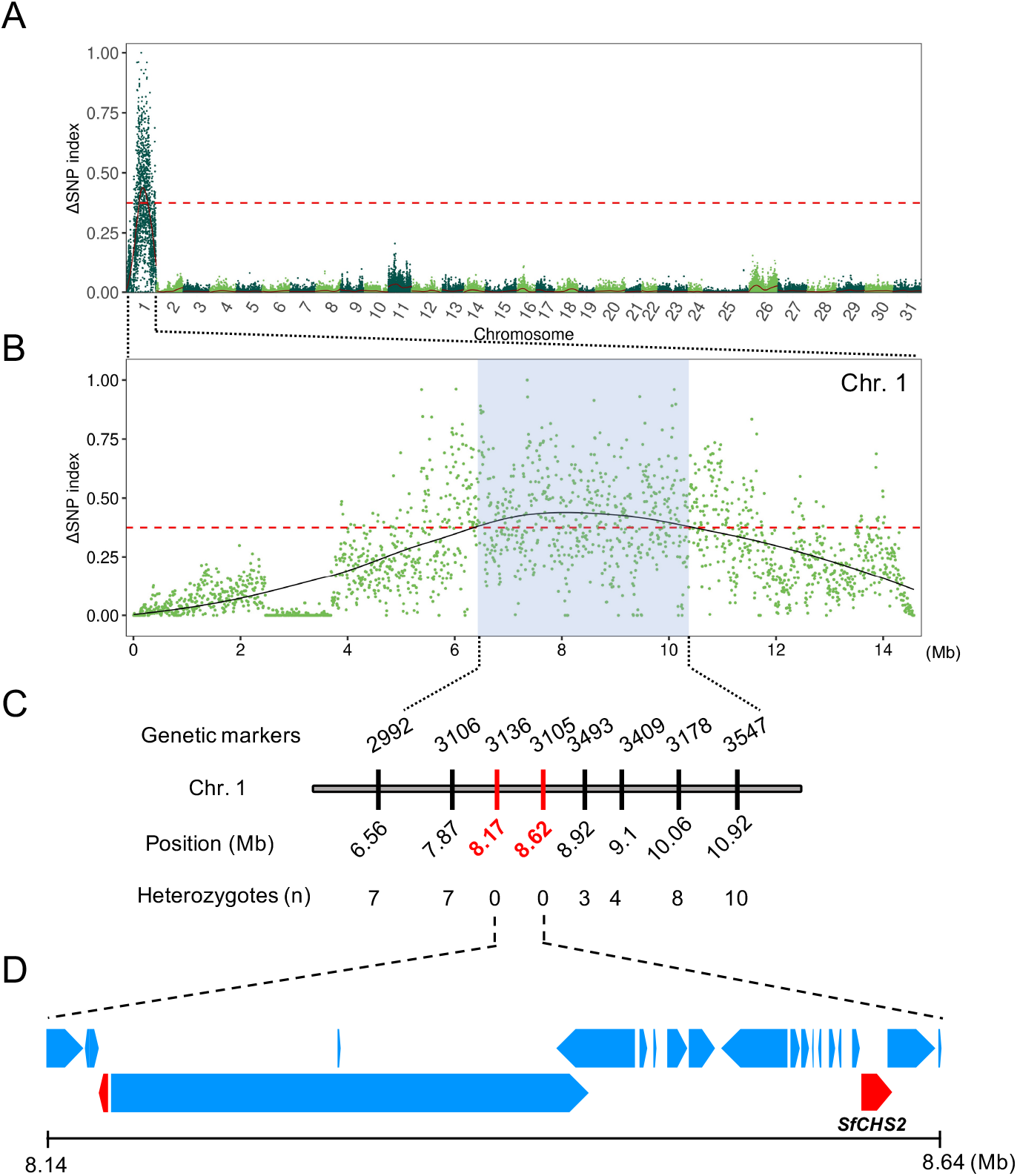
GWAS mapping of the Vip3Aa resistance region. (A) Manhattan plot for bulked segregant analysis. The raw reads were filtered using trimmomatic, then clean reads were mapped onto *Spodoptera frugiperda* reference genome AGI-APGP_CSIRO_Sfru_2.0 (GCF_023101765.2). GATK was used to perform variant calling, such as Single Nucleotide Polymorphisms (SNPs) and InDels. The association analysis was comprehensively evaluated using SNP-index. The top 1% Δ (SNP-index) region was identified as candidate region. (B) Chromosome 1 showing a maximum of the ΔSNP-index in a 4.36 Mb interval (blue). (C) Eight markers in the 4.36 Mb interval. Only the last 4 digits of the marker ID are shown; e.g. the full name for the first one is LOC118272992. (D) 16 genes in the smaller 0.45 Mb interval, with *SfCHS2*, chitin synthase 2 in red on the right.

A total of eight genetic markers (2992, 3106, 3136, 3105, 3493, 3409, 3178 and 3547) distributed uniformly on chromosome 1 (6.56, 7.87, 8.17, 8.62, 8.92, 9.1, 10.06 and 10.92 Mb) were used to further narrow the genetic interval linked with Vip3Aa resistance (Figure 1C). Two of the eight genetic markers, 3136 and 3105, were shown to be tightly linked with Vip3Aa resistance, since all the tested F7-R progeny were homozygotes for the alleles derived from Sfru_R3 (Figure 1C). In contrast, for the other genetic markers, heterozygotes were detected in tested F7-R progeny and the number of heterozygotes gradually increased with the distance away from marker 3136 or 3105 (Figure 1C). Subsequently, we further focused on those genes included in the genomic interval ranging from 8.17 to 8.62 Mb in Chr.1.

In the narrowed genomic interval, a total of 18 genes were annotated and most of them were transcribed at a relatively low level or not at all, in the midguts of larvae from SS (Table S6). Specifically, RNA-seq analysis showed that LOC118273137, LOC118273449 and LOC118273105 exhibited relative higher transcription level in the midgut tissue compared with the other genes in the candidate interval (Table S6). Structural analysis of predicted proteins encoded by the 18 genes showed that only LOC118273138 and LOC118273105 were able to produce plasma membrane-located protein. However, the FPKM (Fragments Per Kilobase of transcript per Million mapped reads) value of LOC118273138 in midgut was merely 0.28, indicating a likely lack of transcription for this gene in the midgut tissue (Table S6). Taken together the transcriptional quantity in midgut and predicted protein structure, we selected LOC118273105 as the primary candidate gene for further investigation.

### A LTR retrotransposon insertion results in the decrease of wild-type *SfCHS2* transcripts

The LOC118273105 was annotated as chitin synthase (CHS), a membrane-integral glycosyltransferase, which catalyzes chitin biosynthesis via transferring GlcNAc from UDP-GlcNAc to a growing chitin chain. Another CHS gene, LOC118273150, located next to LOC118273105 was annotated as well. Further phylogenetic analysis showed that LOC118273105 and LOC118273150 were identified as the *CHS2* and *CHS1* in *S. frugiperda*, respectively (Fig. S2). Thus, we named LOC118273105 and LOC118273150 as *SfCHS2* and *SfCHS1*, respectively. Results from RNA-Seq analysis demonstrated that *SfCHS2* was transcribed at a high level in the larvae midgut tissue of *S. frugiperda*, while *SfCHS1* was transcribed at a low level (Table S6), suggesting a more potential role of *SfCHS2* than *SfCHS1* in the mode of action of Vip3Aa toxin.

To investigate the potential InDel mutation in *SfCHS2*, the full length genomic sequences of *SfCHS2* in SS and Sfru_R3 were amplified by PCR. Results from Sanger sequencing showed that a large fragment (6432 bp) insertion between exon21 and intron21 of *SfCHS2* was observed in Sfru_R3 (Figure 2A). Interestingly, BLAST results demonstrated that the inserted sequence was a long terminal repeat (LTR) retrotransposon (“Yaoer”, 幺蛾, wicked), which was composed of two LTRs (LTRa and LTRb) at the 5’ and 3’ terminal and a protein encoding sequence that contains the aspartate proteases (AP), reverse transcriptase (RT), ribonuclease H (RNase H) and integrase (INT) domains (Figure 2B). The insertion point was identified according to the target site duplication (TSD) on both ends of the retrotransposon. The sequence GAAGG was identified immediately before the start of LTRa and immediately after the end of LTRb (Fig. S3). Although the LTR retrotransposon was inserted between exon21 and exon22, the original 5’ splice site of the intron21 is intact, and completely to the right of the transposable element (Fig. S3). Basically the transposable element has pushed the intron further downstream on the genomic DNA. Since all of the necessary sequences of intron21 are present (5’ GU, internal branch point, 3’AG), it could be spliced out as usual, retaining the LTR retrotransposon following the exon21. However, the possibility that LTR retrotransposon was spliced out completely along with intron21 cannot be readily excluded, since a new 5’ splice site (GU) was observed following the 3’ terminal of exon21 (Fig. S3). The latter splicing would result in a truncated protein partially identical (initial 1343 of 1524 residues) with the wild-type *SfCHS2* protein (Fig. S4).

**Figure 2.**
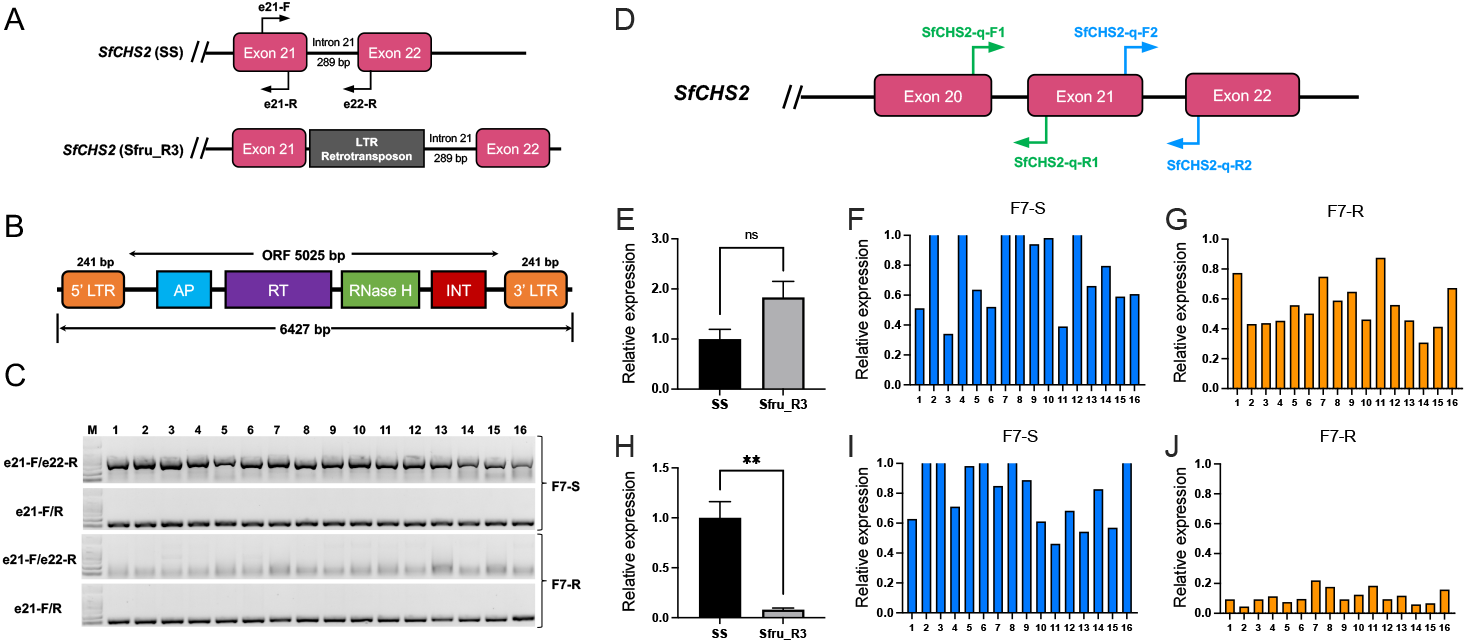
LTR retrotransposon insertion causes reduced wild-type *SfCHS2* transcripts in Sfru_R3. (A) Comparison of genomic sequences between the 21th intron in Vip3Aa-susceptible (SS) and resistant (Sfru_R3) *S. frugiperda*. The red box, black horizontal line and grey box refers to exon, intron and retrotransposon, respectively. (B) Diagram of the LTR retrotransposon insertion. The LTR retrotransposon harbors two LTR region at the 5’ and 3’ terminal, respectively. The protein region contains aspartate proteases (AP), reverse transcriptase (RT), ribonuclease H (RNase H) and integrase (INT) domains. (C) Genotype identification of *SfCHS2* from Vip3Aa-susceptible and resistant larvae in F7 population based on PCR with genomic DNA. A pair of primers (e21-F/e22-R) flanking the 21th intron were used and their locations were indicated as Fig. 2A. Parallelly, primers pair (e21-F/e21-R) in exon21 were used as the positive control. The LTR insertion was genetically linked with resistance to Vip3Aa toxin (*P* < 0.01, *χ*^2^ test). (D) The locations of primers used for relative expression investigation of *SfCHS2* via RT-qPCR. A pair of primers (*SfCHS2*-q-F1/R1) that located at the 20th and 21th exon were designed to evaluate the relative expression level of total transcripts of *SfCHS2* in Vip3Aa-susceptible and resistant strain (E), F7-S (F) and F7-R (G) individuals. Another pair of primers (*SfCHS2*-q-F2/R2) flanking the 21th intron were designed to test the wild-type transcripts of *SfCHS2* in Vip3Aa-susceptible and resistant strain (H), F7-S (I) and F7-R (J) individuals. The relative higher expression of total *SfCHS2* transcripts observed in Sfru_R3 was not genetically linked with resistance to Vip3Aa (*P* > 0.05, *χ*^2^ test). In contrast, the reduced wild-type *SfCHS2* transcripts was genetically linked with resistance to Vip3Aa in Sfru_R3 (*P* < 0.01, *χ*^2^ test). The relative expression level of *SfCHS2* in Sfru_R3 and individuals from F7-S and F7-R were normalized to the fold value of 2^-ΔCt^ relative to that in SS. The double asterisk “**” refers to *P* < 0.01(*t*-test, df=3). Ns: not significant.

To test whether the LTR retrotransposon detected in Sfru_R3 was spliced out or not, we firstly investigated the transcripts of *SfCHS2* using a pair of primers located at exon21 and exon22. Results from gel electrophoresis and Sanger sequencing showed that the transcripts without Yaoer were able to be amplified in Sfru_R3 (Fig. S5A), suggesting the complete splice out of Yaoer and intron21. However, we also detected the transcripts that retain the Yaoer in Sfru_R3 using a pair of primers located at the exon21 and 5’ terminal region of LTR (Fig. S5A). These results demonstrate that both wild-type and Yaoer-included transcripts of *SfCHS2* were produced simultaneously in Sfru_R3. Interestingly, the transcripts including intron21 were also detected in Sfru_R3 (Fig. S5A), suggesting potential non-splicing transcripts, including Yaoer and intron21 simultaneously, might be generated by the Yaoer insertion event. In addition, results from ISO-seq indicate that both wild-type and mutant *SfCHS2* transcripts with Yaoer were produced in Sfru_R3 (Fig. S5B).

Based on those results, we speculated that the relative expression level of wild-type *SfCHS2* transcripts in Sfru_R3 might be affected and different with that in SS. As we predicted, no significant (*t*-test, *P* = 0.0776) difference in total transcripts of *SfCHS2* was observed between Sfru_R3 and SS (Figure 2E). Interestingly, however, significant decrease (*t*-test, *P* = 0.0032) of wild-type *SfCHS2* transcripts was detected in Sfru_R3 (0.08 ± 0.016, df = 3) compared with SS (1.0 ± 0.162, df = 3) (Figure 2H). Thus, we speculated that the reduction of wild-type *SfCHS2* transcripts could be associated with resistance to Vip3Aa in Sfru_R3.

### Causal relationship between reduction of wild-type *SfCHS2* transcripts and Vip3Aa resistance in Sfru_R3

To test the potential causality between reduction of the wild-type *SfCHS2* transcripts and resistance to Vip3Aa in Sfru_R3, we firstly investigated the relevance of Vip3Aa resistance with Yaoer insertion. Results from genetic linkage analysis demonstrated that all survivors from F7 progeny (F7-R) were the homozygotes of resistance allele of *SfCHS2* with Yaoer insertion (Figure 2C). In contrast, all tested susceptible individuals from F7 progeny (F7-S) after exposure of Vip3Aa toxin harbored the wild-type allele without Yaoer insertion (Figure 2C). These results suggest that the Yaoer insertion was genetically linked with resistance to Vip3Aa in Sfru_R3 (*P* < 0.01, *χ*^2^ test). Then, we investigated the relative expression level of *SfCHS2* transcripts in the midgut from SS, Sfru_R3, F7-S and F7-R larvae. Results from RT-qPCR analysis showed that all tested F7-R larvae exhibited a significant suppression (0.114 ± 0.012, *n* = 16) in the wild-type *SfCHS2* transcripts, while all tested F7-S larvae exhibited normal expression level (1.159 ± 0.205, *n* = 16) of wild-type *SfCHS2* transcripts that was comparable with that in SS (Figure 2I and J), suggesting the reduction of wild-type *SfCHS2* transcripts was genetically linked with resistance to Vip3Aa in Sfru_R3 (*P* < 0.01, *χ*^2^ test). However, the relative expression level of total *SfCHS2* transcripts in Sfru_R3 was not genetically linked with Vip3Aa resistance (*P* > 0.05, *χ*^2^ test) (Figure 2F and G). Based on that, we further proposed that reduction of wild-type *SfCHS2* transcripts, caused by Yaoer insertion, could be responsible for high-level resistance to Vip3Aa in Sfru_R3.

Subsequently, we established a *SfCHS2*-knockout strain (SfCHS2-KO-A) of *S. frugiperda* via the CRISPR/Cas9 system, to confirm the causality between reduced expression of wild-type *SfCHS2* transcripts and resistance to Vip3Aa. A 1002 bp deletion between exon4 and exon5 in the *SfCHS2* genomic sequence was generated via a double-sgRNAs knockout strategy (Figure 3A). The large fragment deletion was predicted to result in a truncated protein of *SfCHS2*, in which only the initial 247 amino acids of *SfCHS2* were retained. Results from bioassay with Vip3Aa protoxin showed that SfCHS2-KO-A exhibited a >12,307 resistance ratio relative to SS (Table S7), indicating complete resistance to Vip3Aa (Figure 3B). Results from the complementary test showed that F1 progeny from reciprocal crosses of Sfru_R3 and SfCHS2-KO-A also exhibited similar high-level resistance to Vip3Aa toxin with that in SfCHS2-KO-A (Table S7). To further confirm the causal relationship between *SfCHS2* and Vip3Aa resistance in *S. frugiperda*, another two *SfCHS2*-knockout strains (SfCHS2-KO-B and SfCHS2-KO-C) were established using a double-sgRNAs strategy as well (Fig. S6). There were 86 and 89 amino acids, identical with the wild-type SfCHS2 protein, retained in SfCHS2-KO-B and SfCHS2-KO-C, respectively. Consistent with SfCHS2-KO-A, both SfCHS2-KO-B and SfCHS2-KO-C exhibited complete resistance to Vip3Aa (Table S7). These results together indicate that dysfunction of *SfCHS2* is able to cause complete resistance to Vip3Aa in *S. frugiperda* and that genetic element underlying the resistance to Vip3Aa in Sfru_R3 is located at the same locus as *SfCHS2*.

**Figure 3.**
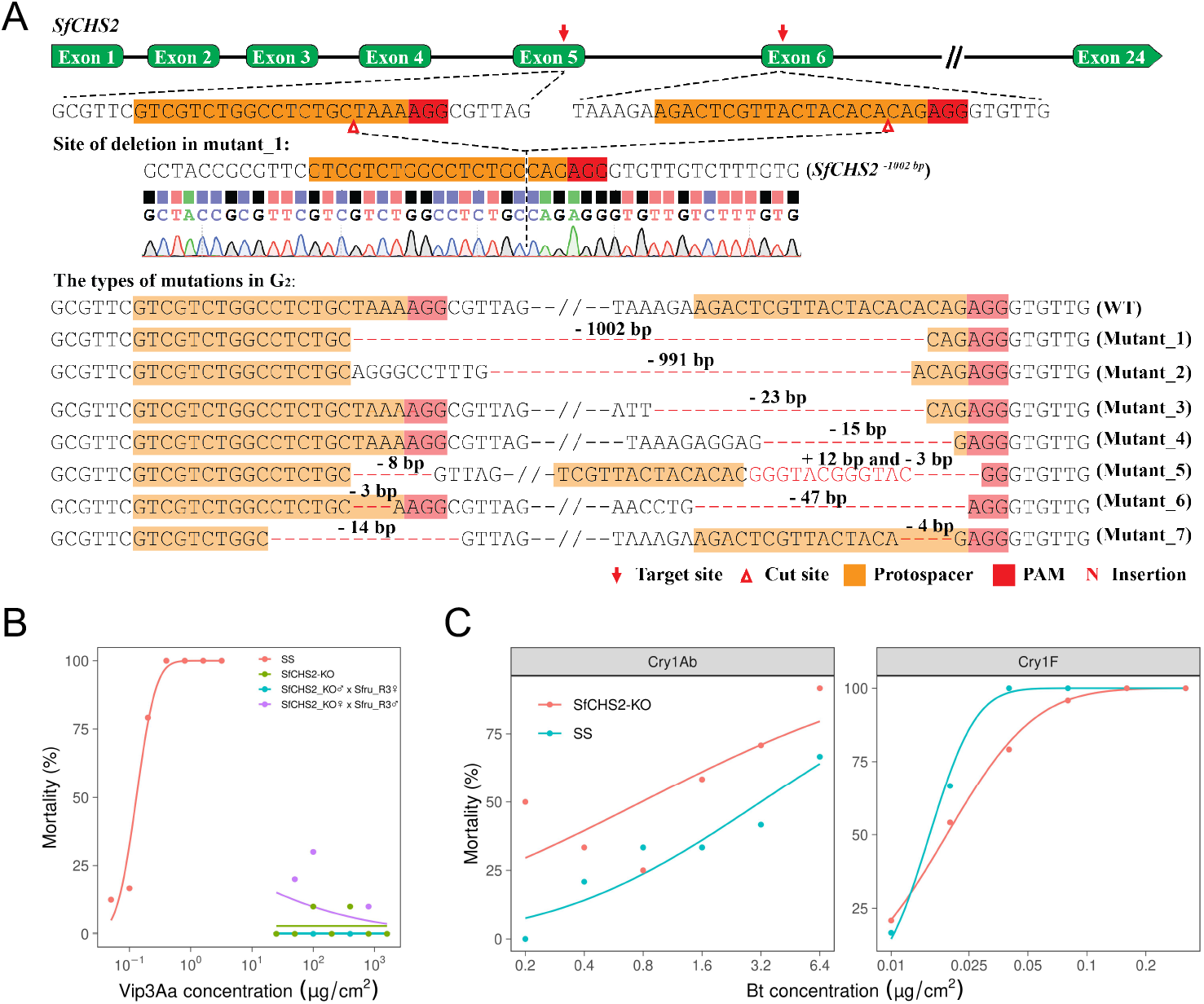
Knockout of *SfCHS2* via CRISPR/Cas9 causes complete resistance to Vip3Aa in *S. frugiperda*. (A) CRISPR/Cas9-mediated double sgRNA system and various types of mutations in G1 larvae identified through sequencing of individual PCR clones. Deleted bases are indicated as red dashes, and inserted bases are indicated as red letters. The CRISPR target sites and the number of deleted and inserted bases (+, insertion; –, deletion) are shown. The chromatogram shows the sequence of the mutant isolated from a homozygous knockout larva in G2. (B) Log dose–response curves for the SS, SfCHS2-KO and the progeny of reciprocal crosses from SfCHS2-KO and Sfru_R3 strains exposed to Vip3Aa protoxin. Both SfCHS2-KO and the progeny of crosses with Sfru_R3 exhibited complete resistance to Vip3Aa. The defining feature of complete resistance is a flat mortality-concentration curve or effect-concentration curve (zero slope) over a wide range of high toxin concentrations. (C) Absence of cross resistance to other Bt toxins in SfCHS2-KO. No cross-resistance to Cry1Ab and Cry1F toxin were observed in the SfCHS2-KO strain.

### Reduced thickness of peritrophic matrix in Vip3Aa-resistant *S. frugiperda* larvae

To test whether reduction of wild-type *SfCHS2* transcripts could affect the peritrophic matrix (PM) structure or not, we dissected the midgut and compared the thickness of PM in larvae from SS, Sfru_R3 and SfCHS2-KO. From the full view of midgut, we observed that gut contents of SS and Sfru_R3 are enclosed by a glossy transparent peritrophic matrix with no evident leakage of Blue Dextran (Fig. 4A). There is no obvious difference in the midgut structure, including PM, between SS and Sfru_R3. In contrast, the SfCHS2-KO has no apparent PM, and the food bolus retained the torpedo-like shape for only 60 seconds before collapsing into a heap after no longer being contained in the midgut (Fig. 4A). However, after cutting terminal of midgut and fixed with picric acid, we observed that the transparent PM in Sfru_R3 was shorter and thinner than that in SS (Fig. 4B). Furthermore, results from histological structure analysis with transverse section through midgut showed that the PM in Sfru_R3 was thinner than that in SS (Fig. 4C). Also, no obvious PM structure was observed in SfCHS2-KO (Fig. 4C). These results together suggest that reduction of wild-type *SfCHS2* transcripts may result in, via less chitin synthesis, the decrease of thickness of PM in Sfru_R3.

**Figure 4.**
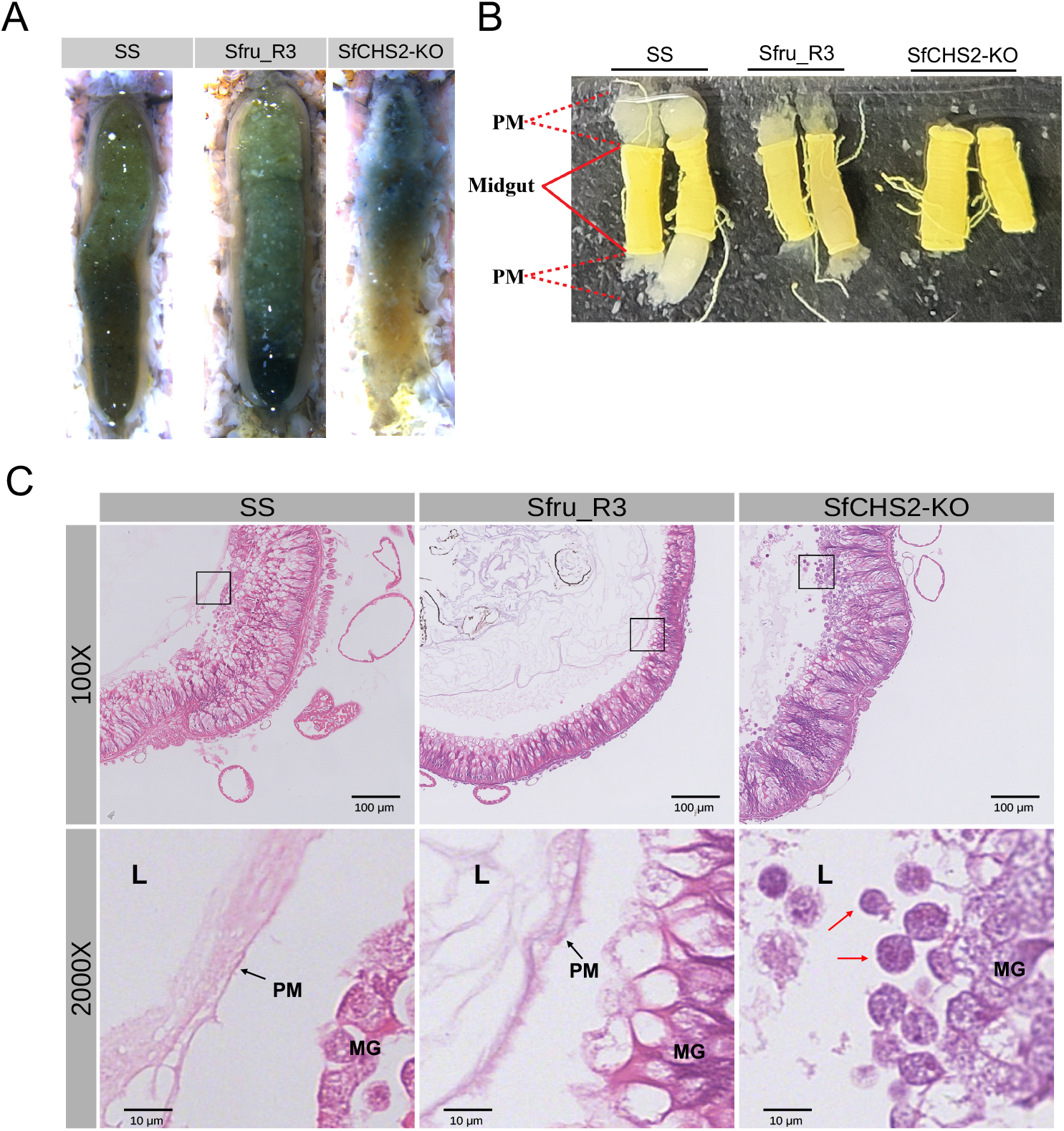
Dissected midguts and peritrophic matrix in SS, Sfru_R3 and *SfCHS2*-KO larvae of *S. frugiperda*. (A) Fifth-instar larvae feeding on artificial diet were starved for 4 hours then allowed to imbibe a sucrose solution containing Blue Dextran 2000 and transferred back to diet. Midguts were removed, a longitudinal incision made and the cut midgut edges pinned back to display the contents. Gut contents of SS and Sfru_R3 are enclosed by a glossy transparent peritrophic matrix with no evident leakage of Blue Dextran. The SfCHS2-KO has no apparent peritrophic matrix, and the food bolus retained the torpedo-like shape for only 60 seconds before collapsing into a heap after no longer being contained in the midgut. (B) Midguts after fixation with picric acid. (C) Histological structure of transverse section through midgut. The peritrophic matrix was thick and compact in SS strain. But it was relatively thin and loose in Sfru_R3 strain. There was no peritrophic matrix, but a lot of shed cells in lumen of midgut in SfCHS2-KO strain (red arrowhead). L, lumen of midgut. MG, midgut. PM, peritrophic matrix.

### Knockout of *CHS2* results in complete resistance to Vip3Aa toxin in *Spodoptera litura* and *Mythimna separata*

To investigate whether *CHS2* also plays a common role in the other species, we established *CHS2*-knockout strains of *S. litura* and *M. separata* and evaluated their susceptibilities to Vip3Aa toxin. For the SlCHS2-KO strain, the mutation of *SlCHS2* was a 207 bp deletion between exon3 and exon 4 (Fig. S7A), For the MsCHS2-KO strain, the mutation of *MsCHS2* was a 366 bp of deletion between exon4 and exon5 (Fig. S7B). Consistent with SfCHS2-KO strains, results from bioassays showed that SlCHS2-KO and MsCHS2-KO exhibited more than 100,000 and 1,333-fold resistance ratio relative to corresponding susceptible strain, respectively (Table S8). These results indicate that *CHS2* also involves in facilitating the toxicity of Vip3Aa in *S. litura* and *M. separata*.

### Detection of Yaoer insertion in *SfCHS2* from a field population of *S. frugiperda*

To determine whether the Yaoer insertion in *SfCHS2* was evolved in the field or not, the genomic reads of a total of 540 samples collected from the field or laboratory were mapped with the junction of exon21 or exon22 and Yaoer (33, 34). Interestingly, one sample (Sample ID: GDJM14) from Jiangmen city, Guangdong province (China), in 2020 showed positive mapping reads (Fig. 5). In this particular sample, the left and right insertion sites were covered by four and six reads, respectively (Fig. S8A). Notably, no reads covered the exon21-intron21 junction site. In contrast, reads from the remaining 539 samples covered the exon21-intron21 junction site only (Fig. S8B). The detection of the Yaoer insertion in insects from the field suggests that this insertion event may has occurred before the selection process in laboratory.

**Figure 5.**
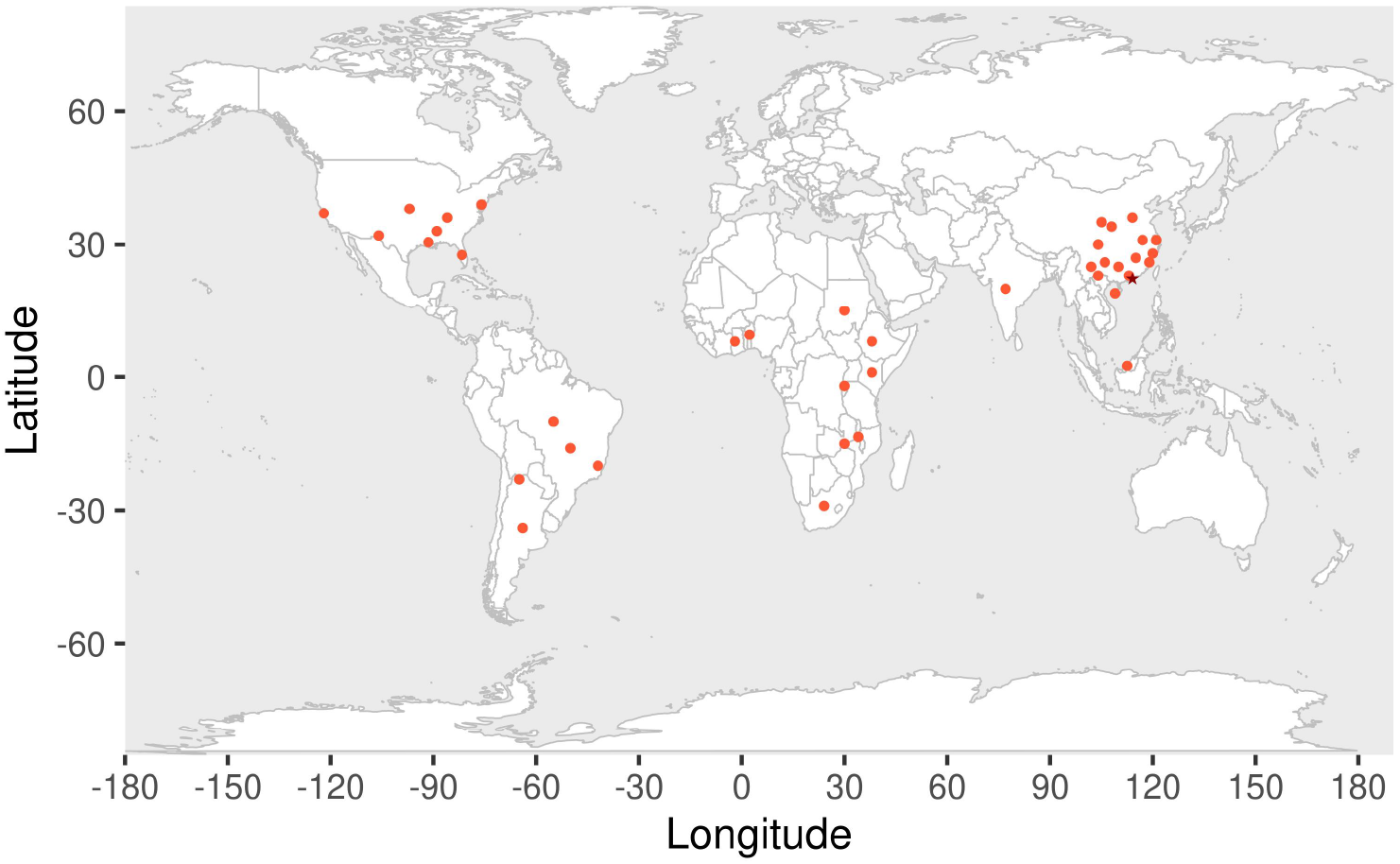
Detection of Yaoer insertion in *SfCHS2* from the field and laboratory population of *S. frugiperda* around the world. A total of 540 *S. frugiperda* individuals collected from the field or laboratory around the world were re-sequenced and mapped with the junctions of exon21 or intron21 and Yaoer element. Among those, one individual collected from the field of China in 2020 showed positive mapping. The red dots refer to locations for all collected samples. The red star refers to location for the positive mapping individual. The detailed information for all tested individuals were summarized as (33, 34).

## Discussion

In the current study, we report a highly Vip3Aa-resistant population of *S. frugiperda* selected in laboratory. All results from BSA, genetic linkage analysis and CRISPR/Cas9 gene editing support that the midgut-specific chitin synthase, *SfCHS2*, was associated with high-levels resistance to Vip3Aa in *S. frugiperda*. Specifically, an intact LTR retrotransposon insertion in *SfCHS2* was identified to cause the reduction of wild-type *SfCHS2* transcripts and further lead to high-levels resistance to Vip3Aa. The genetic markers based on Yaoer insertion in *SfCHS2* could be designed and used to monitor potential heterozygotes harboring this resistance allele in the field.

Interestingly, we detected the same mutant *SfCHS2* allele with Yaoer insertion in one sample collected from the field in China in 2020, suggesting the Yaoer insertion event may have occurred in the field, even before the invasion of *S. frugiperda* to China. Considering very limited time after the invasion of *S. frugiperda* to China since early 2019 and the absence of Vip3Aa selection in the field in China, we pose the question: where is the original source of the resistance allele of *SfCHS2* identified in Sfru_R3? Since 2016, Vip3Aa resistance in *S. frugiperda* has been reported successively in regions of United States and Brazil, where Vip3Aa-transgenic corn and cotton have been cultivated for multiple years (13, 19, 20, 32). The widely accepted west-to-east spread hypothesis of invasive *S. frugiperda* across the world supports that the resistance allele of *SfCHS2* likely originated from the United States or Brazil, in which the resistance allele has evolved and distributed widely under selection stress from Vip3Aa-transgenic crops (22, 24). Yet we haven’t detected the Yaoer insertion in all tested samples collected from those fields in which Vip3Aa-transgenic crops have been planted. It may attributed to the relatively limited sample size tested in this study. Thus, to verify this speculation, it is urgently needed to extensively investigate the allele frequency of the mutant *SfCHS2* allele with Yaoer insertion in those fields.

Although scavenger receptor C (SR-C) and fibroblast growth factor receptor (FGFR) have been suggested to be potential receptors for Vip3Aa toxin and supposed to be associated with Vip3Aa resistance (27, 28), resistance alleles of both genes have not been reported in either laboratory- or field-evolved resistant populations so far. Furthermore, knockout of either SR-C or FGFR via CRISPR/Cas9 does not cause significant susceptibility alteration in *S. frugiperda* (35). The downregulation of *SfMyb* causes just 206-fold resistance to Vip3Aa in a laboratory-selected strain, in which the transcripts of *SfCHS2* have not been affected (31). However, in this study, knockout of *SfCHS2* in *S. frugiperda* was demonstrated to confer complete resistance to Vip3Aa toxin. Moreover, reduction of wild-type *SfCHS2* transcripts can also cause remarkably high-levels of resistance (5,562-fold resistance ratio). Those results together with the findings in this study suggest that susceptibility of *S. frugiperda* to Vip3Aa toxin are likely determined by multiple factors, while mutations in *SfCHS2* have more potential to cause high-levels of resistance. Based on the current findings, it is worthwhile to test whether *SfCHS2* is altered or not in those previously reported highly Vip3Aa-resistant populations of *S. frugiperda* derived from the field or selected in laboratory.

From an evolutionary perspective, the variation of *SfCHS2* observed in Sfru_R3 might be the most suitable option for adaptation to Vip3Aa stress. In that case, the high level of resistance was generated, while some wild-type *SfCHS2* transcripts were preserved for chitin synthesis, thus reducing the fitness costs and enhancing the competitiveness of the mutant population. As we predicted, the almost intact PM structure observed in larval midgut from Sfru_R3 supports that residual *SfCHS2* activity was able to produce enough chitin to maintain the normal PM structure (Fig. 5A). There is evidence that the presence of chitin is necessary to maintain PM barrier function in insects (36, 37). When *CHS2* was suppressed, the growth of *Tribolium castaneum* larvae decreased significantly (38). However, the knockdown of *SlCHS2* had no significant effect on the phenotype of *S. litura* (39). Some results support that host plants may amplify the adaptation cost by hindering feeding performance (40). However, the identical relative survival of Sfru_R3 compared with SS when feeding non-Bt maize suggest almost no fitness cost was accompanied with the Yaoer insertion in *SfCHS2* (Table S2). These results imply that the resistance allele of *SfCHS2* with Yaoer insertion has great potential to spread in the field. Indeed, we have detected the same *SfCHS2* allele with Yaoer insertion in a sample collected from the field in China in 2020. This could impact the durability of Vip3Aa-expressing crops.

Another key issues that remains to be further addressed is the specific role of *CHS2* plays in the mode of action of Vip3Aa toxin. Similar with the confirmed receptors for Cry proteins, ATP-binding cassette transporter C2 (ABCC2), CHS2 is a membrane-bound protein that contains multiple transmembrane domains and is highly expressed in the midgut tissue. In addition, immunohistochemistry and immunofluorescence analysis showed that *CHS2* protein was located at the midgut brush border membranes and apical ends of microvilli in *Manduca sexta* (41).

Moreover, knockout of *SfCHS2* can confers *S. frugiperda* complete resistance to Vip3Aa. These features of *CHS2* are similar to those of ABCC2, which has been confirmed to be the primary receptor for Cry1Ac toxin and facilitate pore formation. Thus, *CHS2* is very likely the receptor for Vip3Aa toxin in the midgut tissue of target pests. The most convincing evidence would be results from cytotoxicity tests with cells over-expressing *CHS2*. However, as far as we know, there is no report about the CHS2 overexpression in insect or mammalian cell line. A CHS from fungi, *PsCHS1*, in contrast, has been successfully overexpressed in mammalian HEK293f cells (42).

Here, however, our results from heterologous expression showed that *SfCHS2* was mostly located in the cytoplasm, rather than the predicted plasma membrane of all the three tested cells (mammalian HEK293T, insect Hi5 and Sf9 cells) (Fig. S9A). Furthermore, the cytotoxicity of Vip3Aa to HEK293T and Hi5 cells was not enhanced by overexpression of SfCHS2 (Fig. S9B). This all suggests that correct subcellular location of *SfCHS2* in mammalian or insect cells could be more difficult than fungi CHS. Thus, to clarify the role of *CHS2* in mode of action of Vip3Aa toxin, correct location of CHS2 at the plasma membrane in insect or mammalian cell might be the primary challenge needing to be addressed in the next work.

On the basis of results from PCR, possibly two mutant transcripts were produced by the resistance allele of *SfCHS2* in Sfru_R3, as well as wild-type *SfCHS2* transcripts. Both mutant transcripts were predicted to produce the identical truncated protein of SfCHS2, in which an extracellular structure, C7 domain, was predicted to be lost (43). The association between resistance to Vip3Aa and reduction of wild-type *SfCHS2* transcripts suggest that C7 domain could be a key structure for the toxicity of Vip3Aa mediated by *SfCHS2*. Thus, we propose that the C7 domain at the C-terminus of SfCHS2 is a binding site for Vip3Aa toxin, or for an endogenous protein that interacts with the toxin, or both. A chymotrypsin-like serine protease (GenBank Accession No. XM_035581989.2) homologous to *M. sexta* CLTP1 which binds to C7 could be involved in cleavage of the protoxin and activation of Vip3Aa (43). Alternatively, Vip3Aa itself could bind directly to C7, by means of Domains II and III but probably not the chitin-binding Domain V (44).

In addition, according to its original function in chitin synthesis, the possibility that CHS2 participates indirectly in the toxicity activation of Vip3Aa toxin cannot be excluded. A recent study demonstrated that the binding between Domain V of Vip3Aa toxin and the PM via glycan-binding activity can contributes to Vip3Aa insecticidal activity (44). Chitin is one of the major structural components for PM (45). Thus, it is tempting to speculate a potential association between chitin content in PM and susceptibility of insects to Vip3Aa toxin. However, the variation of susceptibility to Vip3A among lepidopteran insects does not support that speculation (10, 46). On the other hand, that speculation is contrary to the cytotoxicity of Vip3A against Sf9 and Sf21 cells, in which no PM structure was formed (27, 28). Moreover, the complete absence of PM in our CRISPR/Cas9 mutant SfCHS2-KO is correlated with complete resistance to Vip3Aa. Yet the Yaoer insertion does not eliminate the PM in the Sfru_R3 strain, while conferring very high resistance to Vip3Aa. Although the PM in Sfru_R3 larvae appears to be thinner than that in SS, this is unlikely to result in the remarkably high-levels resistance to Vip3Aa, compared with the complete absence of PM in SfCHS2-KO larvae. Thus, the presence of a PM appears to be necessary, but not sufficient for the toxicity of Vip3Aa to larvae.

Unexpectedly, even though the PM structure in SfCHS2-KO was shown to be completely invisible, and the midgut was directly exposed under the ingested food, insects of the knockout lines complete the larval stage on artificial diet and pupate (Table S9), and adults are fertile. Although the PM confers benefits in protecting the midgut epithelium from abrasion, viral infection, and bacteria in nature (47), it is not required for digestion and nutrient assimilation when feeding artificial diet. Our results provide a new perspective on chitin synthesis by completely eliminating the PM for the first time, by deleting chitin synthase 2.

Through investigations conducted on *S. litura* and *M. separata*, closely and distantly related species of *S. frugiperda*, it was found that the disruption of the CHS2 gene yielded complete resistance to Vip3Aa toxin. These findings indicate that CHS2 could potentially serve as a receptor for Vip3Aa toxin within Lepidoptera and potentially even in more distantly related insect taxa, where it could be useful for resistance monitoring.

Transposable elements (TEs) are considered as powerful drivers for genome evolution. In insects, TEs are involved in aging, antiviral immunity and adaptations, such as insecticide resistance (48). For Bt resistance, multiple TE insertion events have been characterized in the receptors for Cry1Ac toxin (49-53). Most of the TE insertion events occurred in exons sequence and result in the production of truncated receptor proteins for Bt toxins. A recent study identified a short interspersed nuclear element (SINE) retrotransposon insertion in the 5’ UTR of *MAP4K4*, which regulates the expression of multiple receptors for Cry1Ac toxin in *Plutella xylostella* (54). Here, we found that a LTR insertion event, for the first time, was associated with pest resistance to Vip3Aa toxin. These results together with previous studies reflect the key role of TEs in insect adaptation to Bt toxins.

Overall, in the current study, we identified a LTR insertion in *SfCHS2* which was associated with the high-level resistance to Vip3Aa in *S. frugiperda*. Furthermore, knockouts of *CHS2* can result in complete resistance to Vip3Aa in *S. frugiperda*, and the other two related species. These results indicate a substantial role that *CHS2* plays in the mode of action of Vip3Aa toxin among lepidopteran species. The first resistance allele of *SfCHS2* identified in this study could not only be applied immediately to monitor Vip3Aa resistance alleles in the field, but also provide a reference for the development of more potential genetic markers for resistance management. The association between *CHS2* and resistance to Vip3Aa demonstrated in this study would significantly boost the elucidation of mode of action of Vip3Aa toxin.

### Materials and Methods

Insect strains, toxin preparation and bioassay, selection of resistance, bioassays on transgenic maize, genetic analysis of resistance, BSA, fine-scale mapping, RNA-Seq, RT-qPCR analysis, genetic linkage analysis, CRISPR/Cas9 knockouts, Yaoer insertion detection in the field, PM structure analysis, overexpression of SfCHS2 in mammalian and insect cells are described in supplemental Materials and Methods.

### Data and materials availability

Sequences of the chitin synthases and the LTR retrotransposon insertion have been deposited in GenBank (Accession Numbers OR669300-OR669302). Code for the BSA analysis is at https://github.com/luming-zou/Yaoer-BSA.

## Acknowledgments

This project was supported by grants from the Sci-Tech Innovation 2030 Agenda (2022ZD04021); National Natural Science Foundation of China (32001944); Innovation Program of Chinese Academy of Agricultural Science (CAAS-CSCB-202303); Agricultural Science and Technology Innovation Program of Chinese Academy of Agricultural Sciences; Major Projects of Basic Research of Science, Technology and Innovation Commission of Shenzhen Municipality; Senior Talents Project of Guangdong (Grant Number 2021A1313030029); and the Max-Planck-Gesellschaft.

## Notes

### Competing Interest Statement

The authors have declared no competing interest.

## References

1. ISAAA, Global status of commercialized biotech/GM crops in 2019: Biotech crops drive socio-economic development and sustainable environment in the new frontier. ISAAA Brief No. 55 (International Service for the Acquisition of Agri-biotech Applications, Ithaca, NY, 2019). (2019).

2. W. D. Hutchison et al., Areawide suppression of European corn borer with Bt maize reaps savings to non-Bt maize growers. Science 330, 222–225 (2010).

3. G. P. Dively et al., Regional pest suppression associated with widespread Bt maize adoption benefits vegetable growers. Proc. Natl. Acad. Sci. U. S. A. 115, 3320–3325 (2018).

4. J. Romeis, S. E. Naranjo, M. Meissle, A. M. Shelton, Genetically engineered crops help support conservation biological control. Biol. Control 130, 136–154 (2019).

5. S. P. Luo, S. E. Naranjo, K. M. Wu, Biological control of cotton pests in China. Biol. Control 68, 6–14 (2014).

6. B. E. Tabashnik, J. A. Fabrick, Y. Carrière, Global patterns of insect resistance to transgenic Bt crops: the first 25 years. J. Econ. Entomol. 116, 297–309 (2023).

7. F. Yang et al., Practical resistance to Cry toxins and efficacy of Vip3Aa in Bt cotton against. Pest Manag. Sci. 78, 5234–5242 (2022).

8. F. Yang, D. L. Kerns, N. S. Little, J. C. Santiago Gonzalez, B. E. Tabashnik, Early warning of resistance to Bt toxin Vip3Aa in Helicoverpa zea. Toxins 13, art. 618 (2021).

9. B. E. Tabashnik, Y. Carrière, Surge in insect resistance to transgenic crops and prospects for sustainability. Nat. Biotechnol. 35, 926–935 (2017).

10. J. J. Estruch et al., Vip3A, a novel Bacillus thuringiensis vegetative insecticidal protein with a wide spectrum of activities against lepidopteran insects. Proc. Natl. Acad. Sci. U. S. A. 93, 5389–5394 (1996).

11. M. K. Lee, F. S. Walters, H. Hart, N. Palekar, J. S. Chen, The mode of action of the Bacillus thuringiensis vegetative insecticidal protein Vip3A differs from that of Cry1Ab delta-endotoxin. Appl. Environ. Microbiol. 69, 4648–4657 (2003).

12. B. R. Pickett, A. Gulzar, J. Ferré, D. J. Wright, Bacillus thuringiensis Vip3Aa toxin resistance in Heliothis virescens (Lepidoptera: Noctuidae). Appl. Environ. Microbiol. 83, e03506–16 (2017).

13. F. Yang et al., F(2) screen, inheritance and cross-resistance of field-derived Vip3A resistance in Spodoptera frugiperda (Lepidoptera: Noctuidae) collected from Louisiana, USA. Pest Manag. Sci. 74, 1769–1778 (2018).

14. B. E. Tabashnik, Y. Carrière, Evaluating cross-resistance between Vip and Cry toxins of Bacillus thuringiensis. J. Econ. Entomol. 113, 553–561 (2020).

15. US-EPA (United States Environmental Protection Agency) (2009) Biopesticide registration action document: Bacillus thuringiensis Vip3Aa20 insecticidal protein and the genetic material necessary for its production via elements of vector PNOV1300 in event MIR162 maize (OECD Unique Identifier: SYN-IR162-4).

16. A. Gulzar, B. Pickett, A. H. Sayyed, D. J. Wright, Effect of temperature on the fitness of a Vip3A resistant population of Heliothis virescens (Lepidoptera: Noctuidae). J. Econ. Entomol. 105, 964–970 (2012).

17. F. Yang, J. C. Santiago Gonzalez, G. A. Sword, D. L. Kerns, Genetic basis of resistance to the Vip3Aa Bt protein in Helicoverpa zea. Pest Manag. Sci. 77, 1530–1535 (2021).

18. F. Yang et al., First documentation of major Vip3Aa resistance alleles in field populations of Helicoverpa zea (Boddie) (Lepidoptera: Noctuidae) in Texas, USA. Sci. Rep. 10, 5867 (2020).

19. X. Chen et al., Fitness costs of Vip3A resistance in Spodoptera frugiperda on different hosts. Pest Manag. Sci. 75, 1074–1080 (2019).

20. O. Bernardi et al., Selection and characterization of resistance to the Vip3Aa20 protein from Bacillus thuringiensis in Spodoptera frugiperda. Pest Manag. Sci. 72, 1794–1802 (2016).

21. R. J. Mahon, S. J. Downes, B. James, Vip3A resistance alleles exist at high levels in Australian targets before release of cotton expressing this toxin. PLoS ONE 7, e39192 (2012).

22. F. S. A. Amaral et al., Geographical distribution of Vip3Aa20 resistance allele frequencies in Spodoptera frugiperda (Lepidoptera: Noctuidae) populations in Brazil. Pest Manag. Sci. 76, 169–178 (2020).

23. J. L. Jurat-Fuentes, D. G. Heckel, J. Ferré, Mechanisms of resistance to insecticidal proteins from Bacillus thuringiensis. Annu. Rev. Entomol. 66, 121–140 (2021).

24. W. T. Tay, R. L. Meagher, Jr., C. Czepak, A. T. Groot, Spodoptera frugiperda: ecology, evolution, and management options of an invasive species. Annu. Rev. Entomol. 68, 299–317 (2023).

25. M. Kenis et al., Invasiveness, biology, ecology, and management of the fall armyworm, Spodoptera frugiperda. Entomol. Gen. 43, 187–241 (2023).

26. X. Hou, L. Han, B. An, J. Cai, Autophagy induced by Vip3Aa has a pro-survival role in Spodoptera frugiperda Sf9 cells. Virulence 12, 509–519 (2021).

27. K. Jiang et al., Scavenger receptor-C acts as a receptor for Bacillus thuringiensis vegetative insecticidal protein Vip3Aa and mediates the internalization of Vip3Aa via endocytosis. PLoS Pathog. 14, e1007347 (2018).

28. K. Jiang et al., Fibroblast growth factor receptor, a novel receptor for vegetative insecticidal protein Vip3Aa. Toxins 10, art. 546 (2018).

29. G. Singh, B. Sachdev, N. Sharma, R. Seth, R. K. Bhatnagar, Interaction of Bacillus thuringiensis vegetative insecticidal protein with ribosomal S2 protein triggers larvicidal activity in Spodoptera frugiperda. Appl. Environ. Microbiol. 76, 7202–7209 (2010).

30. Y. Carrière, B. Degain, G. C. Unnithan, B. E. Tabashnik, Inheritance and fitness cost of laboratory-selected resistance to Vip3Aa in Helicoverpa zea (Lepidoptera: Noctuidae). J. Econ. Entomol. 116, 1804–1811 (2023).

31. M. Jin et al., Downregulation of a transcription factor associated with resistance to Bt toxin Vip3Aa in the invasive fall armyworm. Proc. Natl. Acad. Sci. U. S. A. 120, e2306932120 (2023).

32. Z. M. Wen et al., More than 10 years after commercialization, Vip3A-expressing MIR162 remains highly efficacious in controlling major Lepidopteran maize pests: laboratory resistance selection versus field reality. Pestic. Biochem. Physiol. 192, art. 105385 (2023).

33. L. Zhang et al., Global genomic signature reveals the evolution of fall armyworm in the Eastern hemisphere. Mol. Ecol. 32, 5463–5478 (2023).

34. L. Zhang et al., Genetic structure and insecticide resistance characteristics of fall armyworm populations invading China. Mol. Ecol. Resour. 20, 1682–1696 (2020).

35. Y. Shan et al., Sf-FGFR and Sf-SR-C are not the receptors for Vip3Aa to exert insecticidal toxicity in Spodoptera frugiperda. Insects 13, art. 547 (2022).

36. M. Kelkenberg, J. Odman-Naresh, S. Muthukrishnan, H. Merzendorfer, Chitin is a necessary component to maintain the barrier function of the peritrophic matrix in the insect midgut. Insect Biochem. Mol. Biol. 56, 21–28 (2015).

37. X. Liu et al., Characterization of a midgut-specific chitin synthase gene (LmCHS2) responsible for biosynthesis of chitin of peritrophic matrix in Locusta migratoria. Insect Biochem. Mol. Biol. 42, 902–910 (2012).

38. Y. Arakane et al., The Tribolium chitin synthase genes TcCHS1 and TcCHS2 are specialized for synthesis of epidermal cuticle and midgut peritrophic matrix. Insect Mol. Biol. 14, 453–463 (2005).

39. H. Z. Yu et al., Identification and functional analysis of two chitin synthase genes in the common cutworm, Spodoptera litura. Insects 11, art. 253 (2020).

40. R. H. ffrench-Constant, C. Bass, Does resistance really carry a fitness cost? Curr. Opin. Insect Sci. 21, 39–46 (2017).

41. L. Zimoch, H. Merzendorfer, Immunolocalization of chitin synthase in the tobacco hornworm. Cell Tissue Res. 308, 287–297 (2002).

42. W. Chen et al., Structural basis for directional chitin biosynthesis. Nature 610, 402–408 (2022).

43. G. Broehan, L. Zimoch, A. Wessels, B. Ertas, H. Merzendorfer, A chymotrypsin-like serine protease interacts with the chitin synthase from the midgut of the tobacco hornworm. J. Exp. Biol. 210, 3636–3643 (2007).

44. K. Jiang et al., Functional characterization of Vip3Aa from Bacillus thuringiensis reveals the contributions of specific domains to its insecticidal activity. J. Biol. Chem. 299, 103000 (2023).

45. D. Hegedus, M. Erlandson, C. Gillott, U. Toprak, New insights into peritrophic matrix synthesis, architecture, and function. Annu. Rev. Entomol. 54, 285–302 (2009).

46. C. G. Yu, M. A. Mullins, G. W. Warren, M. G. Koziel, J. J. Estruch, The Bacillus thuringiensis vegetative insecticidal protein Vip3A lyses midgut epithelium cells of susceptible insects. Appl. Environ. Microbiol. 63, 532–536 (1997).

47. M. A. Erlandson, U. Toprak, D. D. Hegedus, Role of the peritrophic matrix in insect-pathogen interactions. J. Insect Physiol. 117, 103894 (2019).

48. C. Gilbert, J. Peccoud, R. Cordaux, Transposable elements and the evolution of insects. Annu. Rev. Entomol. 66, 355–372 (2021).

49. X. Yang et al., Mutation of ABC transporter ABCA2 confers resistance to Bt toxin Cry2Ab in Trichoplusia ni. Insect Biochem. Mol. Biol. 112, 103209 (2019).

50. L. Wang et al., Transposon insertion causes cadherin mis-splicing and confers resistance to Bt cotton in pink bollworm from China. Sci. Rep. 9, 7479 (2019).

51. J. A. Fabrick, L. G. Mathew, B. E. Tabashnik, X. Li, Insertion of an intact CR1 retrotransposon in a cadherin gene linked with Bt resistance in the pink bollworm, Pectinophora gossypiella. Insect Mol. Biol. 20, 651–665 (2011).

52. Y. J. Yang, H. Y. Chen, Y. D. Wu, Y. H. Yang, S. W. Wu, Mutated cadherin alleles from a field population of Helicoverpa armigera confer resistance to Bacillus thuringiensis toxin Cry1Ac. Appl. Environ. Microbiol. 73, 6939–6944 (2007).

53. L. J. Gahan, F. Gould, D. G. Heckel, Identification of a gene associated with Bt resistance in Heliothis virescens. Science 293, 857–860 (2001).

54. Z. Guo et al., Retrotransposon-mediated evolutionary rewiring of a pathogen response orchestrates a resistance phenotype in an insect host. Proc. Natl. Acad. Sci. U. S. A. 120, e2300439120 (2023).

