## Supplemental information for "Retrotransposon-mediated variation of a chitin synthase gene confers insect resistance to *Bacillus thuringiensis* Vip3Aa toxin"

### Supplementary Materials and Methods

#### Insect strains

The SS susceptible strain of *Spodoptera frugiperda* was collected from a maize field in Ruili City, Dehong Prefecture, Yunnan Province, China (24°08′47″N, 97°82′33″E) in 2019 and has been maintained in the absence of any Bt toxins in the laboratory since then. The Vip3Aa-resistant strain, Sfru_R3, was developed from SS by selection on Vip3Aa-incorporated artificial diet. The CRISPR/Cas9-mediated mutant strains SfCHS2-KO-A, SfCHS2-KO-B and SfCHS2-KO-C were constructed starting from SS. Both *Spodoptera litura* and *Mythimna separata* were collected from a field near Beijing in 2017.

#### Toxin preparation and bioassay

The Vip3Aa protoxin was purchased from Beijing Genralpest Biotech Research Co., Ltd (www. genralpest.com), Beijing, China. Vip3Aa protoxin was produced by the BL21 strain of *Escherichia coli*. Vip3Aa protein from cell lysates was purified on a HiTrap Chelating HP column (GE Healthcare, Freiburg, Germany) and was examined for purity by sodium dodecyl sulfate-polyacrylamide gel electrophoresis (SDS-PAGE) analysis. The Vip3Aa protein in the crude cell lysate was quantified by densitometry of the stained band (~88 kDa) on the SDS-PAGE gel. The crude cell lysate was stored at -80 °C until used for bioassays.

Diet overlay bioassays were conducted to assess the toxicity of Vip3Aa protoxin. Artificial diet was poured into 24-well plates and different concentrations of Vip3Aa applied to the surface and allowed to dry. First-instar larvae were transferred to the wells and larval mortality and growth inhibition was assessed after 7 days. Larvae that had not molted to the third instar by 7 days were counted as dead. The $EC_{50}$ (median effective concentration) values and 95% CI (95% confidence interval) were calculated by probit analysis with SPSS 18.0. $EC_{50}$ values with overlapping 95% CIs were considered to be equivalent. Resistance ratios were calculated by dividing the $EC_{50}$ value of each strain by the value of the Vip3Aa-susceptible strain SS.

#### Selection of resistance to Vip3Aa toxin in *S. frugiperda*

Before selection for Vip3Aa resistance, a bioassay test of the susceptible strain was conducted to determine the $EC_{95}$ (effective concentration killing 95% of the larvae). Based on this, the concentration of Vip3Aa for the first round of selection was calculated as 2.22 μg/g. For the first selection, more than 10,000 neonate larvae were inoculated on the artificial diet mixed with Vip3Aa toxin. Five days later, about 400 individuals that developed to the 3rd instar were transferred individually to the normal diet and reared to adults for mating. From the second generation onward, larvae consumed Vip3Aa-treated diet for the entire larval period. The concentration of Vip3Aa was then successively doubled for each of the following three generations. During the selection, 7 bioassays were conducted to determine the real-time level of Vip3Aa resistance to Vip3Aa. At the 16th generation, the concentration of Vip3Aa was increased up to 24 μg/g. A resistance ratio of 5,562 relative to SS was attained at the 17th generation.

#### Bioassays on transgenic maize

We conducted bioassays with different cultivars of maize (*Zea mays*) in a greenhouse at the Chinese Academy of Agricultural Sciences in Beijing. Seeds were obtained from the DBN Group (Beijing) for maize producing Vip3Aa (DBN9501), Cry1Ab (DBN9936), or Cry1Ab + Vip3Aa (DBN9936 × DBN9501), and non-Bt maize (Nonghua106). The concentration of Vip3Aa in DBN9501 was about 25 μg/g by fresh weight. Maize was grown with 5 plants per pot in a greenhouse at about 26 °C. We transferred first instar larvae to maize when the plants were about 30 cm high and had four leaves and one shoot. We measured survival until adult eclosion. For each type of maize and each insect strain (Sfru_R3 and SS), we conducted 3 replicates with 72 larvae per replicate (n = 216 larvae for each combination of maize and insect strain). Measurements were taken from distinct samples; the same sample was not measured repeatedly.

#### Genetic analysis of Vip3Aa resistance in Sfru_R3

Larval mortality in the diet bioassays was calculated as mortality (%) = 100 × (number of dead larvae + number of surviving larvae still in the first instar) / total number of insects assayed. Larval mortality at each Vip3Aa protein concentration was corrected based on the mortality observed on the control diet using the method of Abbott. Probit analysis was used to estimate the $EC_{50}$ and 95% CI as above. In addition, larval mortalities at the Vip3Aa concentrations of 1 - 8 μg/cm$^2$ were analyzed using a one-way ANOVA with insect strain as the main factor. Tests were one-sided.

The dominance of resistance was estimated by comparing the $EC_{50}$s and mortality of the $F_1$ populations with that of the SS and Sfru_R3 parental populations. Dominance levels of Vip3Aa resistance were calculated using two methods. 1) Stone's dominance value *D* compares $EC_{50}$ of different genotypes: $D = (2\log EC_{RS} - \log EC_{RR} - \log EC_{SS}) / (\log EC_{RR} - \log EC_{SS})$, where $EC_{RS}$, $EC_{SS}$ and $EC_{RR}$ are the $EC_{50}$ values of the $F_1$, SS and Sfru_R3 populations, respectively. *D* ranges from -1 (resistance completely recessive) to 1 (resistance completely dominant). 2) The effective dominance $D_{ML}$ compares mortality levels (ML) of different genotypes at a fixed toxin concentration: $D_{ML} = (ML_{RS} - ML_{SS}) / (ML_{RR} - ML_{SS})$, where $ML_{SS}$, $ML_{RR}$, and $ML_{RS}$ are the mortalities of the SS, Sfru_R3, and $F_1$ populations at a given Vip3Aa concentration. $D_{ML}$ ranges from 0 (recessive resistance) to 1 (dominant resistance). Because the mortality and $EC_{50}$ data of the two $F_1$ strains were not significantly different, data from the two $F_1$ strains were pooled to determine the *D* and $D_{ML}$ values.

To determine the inheritance of Vip3Aa resistance in Sfru_R3, the following crosses were conducted: (a) reciprocal parental crosses $F_1$a = Sfru_R3♂ × SS♀ and $F_1$b = Sfru_R3♀ × SS♂ , (b) $F_1$ crosses $F_2$a = $F_1$a × $F_1$a and $F_2$b = $F_1$b × $F_1$b, and (c) Sfru_R3 backcrosses BCR1 = $F_1$a♂ × Sfru_R3♀, BCR2 = $F_1$a♀ × Sfru_R3♂, BCR3 = $F_1$b♂ × Sfru_R3♀, and BCR4 = $F_1$b♀ × Sfru_R3♂.

Chi-squared ($\chi^2$) tests for goodness of fit were used to evaluate whether the inheritance of resistance fitted a Mendelian monogenic model: $\chi^2 = (O - E)^2 / np(1 - p)$, where O is the observed number of dead larvae of the $F_2$ or backcross populations at a certain Vip3Aa concentration, E is the expected number of dead larvae, n is the number of total larvae tested and p is the expected mortality. The null hypothesis was that resistance is controlled by one locus with two alleles: S (susceptible) and R (resistant). If so, the backcross between RS and RR will produce progeny that are 50% RR and 50% RS, the backcross between RS and SS will produce progeny that are 50% SS and 50% RS, and the $F_2$ are expected to consist of 25% RR, 50% RS and 25% SS. The $\chi^2$ value was compared with the $\chi^2$ distribution with 1 degree of freedom, and since *P* was > 0.05, the null hypothesis of monogenic resistance could not be rejected (one-sided test).

**Whole-genome sequencing (WGS) -based bulked segregant analysis (BSA)**

For resistance mapping, a single pair cross (male Sfru_R3 × female SS) was conducted to generate $F_1$ progeny. The head and thorax of the parents were snap-frozen in liquid nitrogen and stored at -80 °C. The $F_1$ progeny were raised on a toxin-free diet, and intercrossed to produce $F_2$ progeny. 480 $F_2$ neonates were placed on a high Vip3Aa diet (4.0 μg/cm$^2$) and 480 $F_2$ neonates were placed on a low Vip3Aa diet (0.1 μg/cm$^2$) for five days. Progeny on the high Vip3Aa diet that molted to the 3rd instar after 5 days (n = 89) were classified as resistant (F2-R). Progeny on the low Vip3Aa diet that had not yet molted to the 3rd instar after 5 days (n = 98) were classified as susceptible (F2-S). These two groups were reared to the 5th instar on toxin-free diet. Each larva was snap-frozen and stored at -80 °C. DNA was isolated separately from Sfru_R3, SS, F2-R (n=50) and F2-S (n=50) samples. Equal amounts of DNA from each individual F2-S larva were mixed, resulting in a bulked (pooled) DNA sample representing the F2-S group; the F2-R group was pooled in a similar manner. The DNA samples were sent to Biomarker (Beijing, China) for whole-genome sequencing on the Illumina NovaSeq 6000 platform. The requested sequence coverage for parents and $F_2$ progenies were 20× and 100×, respectively.

The raw reads were filtered by using trimmomatic, then clean reads were mapped onto the most recent *Spodoptera frugiperda* reference genome at GenBank (AGI-APGP_CSIRO_Sfru_2.0, GCF_023101765.2). Then GATK was used to perform variant calling, including Single Nucleotide

Polymorphisms (SNPs) and InDels. Prior to association analysis, we filtered out SNP sites with multiple genotypes (i.e., more than two genotypes between two extreme traits) and then filtered out SNP sites with read support less than four. Then, we filtered out SNP sites with consistent genotypes between the susceptible ss and resistant rr grandparents and the genotypes of offspring that did not originate from the parents, finally resulting in high-quality SNP loci. Finally, the top 1% Δ (SNP-index) region was identified as the candidate region.

**Fine-scale mapping**

For fine-scale mapping of Vip3Aa resistance, 50 $F_1$ progeny of a SS female × Sfru_R3 male cross were intercrossed to form the $F_2$; 50 $F_2$ progeny were intercrossed to form the $F_3$, and so on for seven generations. Neonates from the $F_7$ generation were divided into two groups: >10,000 larvae on high Vip3Aa diet (24.0 μg/cm$^2$) and 2,400 larvae on low Vip3Aa diet (0.1 μg/cm$^2$) for five days. The 214 largest survivors on the high Vip3Aa diet formed the F7-R group and the 109 slowest-growing survivors on the low Vip3Aa diet formed the F7-S group. Both groups were reared to the fifth instar on toxin-free diet. The genomic DNA of 157 3rd-instar F7-R larvae and the original parents was isolated individually using the DNeasy Blood & Tissue Kit (Qiagen, Germany). The midguts of F7-R (n=16) and F7-S (n=16) were homogenized and stored in TRIzol reagent (Invitrogen, Carlsbad, USA) at -20°C for RNA isolation.

Based on the genome sequence and homozygous SNPs differentiating SS and Sfru_R3, we designed genetic markers in the exons of eight evenly spaced genes in the candidate genomic region. Specific primers (Table S10) for the eight SNP markers were designed and used to investigate the genotypes of 157 F7-R individuals by PCR amplification and sequencing. Denoting SNP alleles from Sfru_R3 as r and those from SS as s, the numbers of rr or ss homozygotes and rs heterozygotes for each locus marker were recorded.

**RNA sequencing (RNA-Seq)**

The midguts of fifth-instar larvae were collected, rapidly frozen in liquid nitrogen, and stored at -80 °C. Total RNA was extracted using the TRIzol reagent according to the manufacturer's instructions. The RNA was sent to BerryGenomics (Beijing, China) for library construction and sequencing. Raw sequence reads were assessed for quality using FastQC. Trimming of adapter sequences and low-quality bases was performed using trimmomatic. The processed reads were aligned to the *S. frugiperda* reference genome (GenBank Accession GCF_023101765.2) using HISAT2. StringTie was employed to assemble transcripts and quantify gene expression levels. StringTie-generated GTF files containing transcript information were processed to create gene-level FPKM tables.

**Genomic DNA isolation, RNA extraction, cDNA synthesis and RT-qPCR analysis**

The gDNA of single fourth-instar larvae was isolated for detection using the AxyPrep Multisource Genomic DNA Miniprep Kit (Axygen, NY, USA) according to the manufacturer’s specifications. PCR reactions were performed in a S1000 Thermal Cycler (Bio-Rad) using the 2×Taq Plus Master Mix II (Vazyme, Nanjing, China) as follows: initial denaturation 95 °C for 3 min, followed by 35 cycles of 95 °C for 15 s, 55°C for 15 s and 72 °C for 30 s, and a final extension of 5 min at 72 °C.

Total RNA was extracted from *S. frugiperda* midgut tissues using TRIzol reagent (Invitrogen) following the manufacturer’s instructions. 1 μg total RNA was used to synthesize the first-strand cDNA using the Revert Aid First Strand cDNA Synthesis Kit (Thermo Fisher Scientific, Waltham, MA, USA). The relative expression levels of targeted genes were quantified by RT-qPCR with AceQ Universal SYBR qPCR Master Mix (Vazyme, Nanjing, China) in a Bio-Rad CFX Connect Real-Time System (Bio-Rad, USA). The reaction was performed in a final volume of 10 µL containing 5 µL of 2×AceQ Universal SYBR qPCR Master Mix, 1 µL of cDNA and 0.5 µL of each primer (10 µM). The reaction conditions were as follows: initial denaturation at 95 °C for 2 min, followed by 40 cycles at 95 °C for 5 s and 60 °C for 30 s. A melting curve analysis was performed after the amplifications to determine the Tm of the amplicons as a quality check. The glyceraldehyde-3-phosphate dehydrogenase gene (GAPDH) was used as an internal control.

Primers were GAPDH_F: 5'-CGG TGT CTT CAC AAC CAC AG-3' and GAPDH_R: 5'-TTG ACA CCA ACG ACG AAC AT-3'. For quantitative analysis of total transcripts of *SfCHS2*, the primers were SfCHS2-q-F1: 5'-TGT TCG TGC TCG TCA TCT TC-3’ and SfCHS2-q-R1: 5'-ACC GAT AGG TTC CAG CGT TA-3’. For quantitative analysis of wild-type transcripts of *SfCHS2*, the primers were SfCHS2-q-F2: 5'-GCC ATG TTG TTC CAT CGC CT-3’ and SfCHS2-q-R2: 5'-AGT CGT CGG TGT TCA GAC GT-3’.Quantitative analysis of gene expression was calculated using the $2^{-\Delta Ct}$ or $2^{-\Delta\Delta Ct}$ method. Primer locations are depicted in Fig. 2D. PCR conditions were initial denaturation 95 °C for 3 min, followed by 35 cycles of 95 °C for 15 s, 55 °C for 15 s and 72 °C for 30 s.

**Detection of *SfCHS2* transcripts via PCR and Isoform-sequencing (ISO-seq)**

For PCR detection, specific primers (*SI Appendix*, Table S10) were designed to detect the potential transcripts in SS and Sfru_R3. The total RNA of midgut tissue from individual larvae were isolated and used to synthesize cDNA as described above. PCR reactions were performed as described above.

For ISO-seq, the total RNA of midguts from 5th-instar larvae of SS and Sfru_R3 were isolated and sent to BerryGenomics (Beijing, China) for library construction and sequencing. The Pacific Biosciences toolkit (https://github.com/PacificBiosciences/pbbioconda) was utilized for ISO-seq data analysis. Initially, raw reads underwent primer removal using lima (v2.7.1) and subsequent refinement involving polyA removal using isoseq3. Following this preprocessing, the clean reads underwent clustering via isoseq3 cluster to generate transcriptional sequences. The isoforms of *SfCHS2* were then extracted using NCBI-BLAST (v2.11.0) and aligned to the reference sequence using Minimap2 (v2.26).

**Genetic linkage analysis**

To test the association between retrotransposon insertion in *SfCHS2* and Vip3Aa resistance, we firstly investigated the genotype of *SfCHS2* in F7-R (n=16) and F7-S group (n=16) using a pair of specific primers (e21-F: 5’-GCC ATG TTG TTC CAT CGC CT-3’/e22-R: 5’-AGT CGT CGG TGT TCA GAC GT-3’) which across the 21th intron and was predicted to amplify a 467 bp fragment of wild-type *SfCHS2* under the following condition: initial denaturation 95 °C for 3 min, followed by 35 cycles of 95 °C for 15 s, 55 °C for 15 s and 72 °C for 30 s. Parallelly, a pair of specific primers (e21-F: 5’-ACC CAA GAC TAC TTA ACG CT-3’/e21-R: 5’-TTT GGT GGT GGA CAG CAG AT-3’) in exon21 were used as the positive control. To further test the genetic association between resistance to Vip3Aa and reduced wild-type transcripts of *SfCHS2*, a pair of specific primers (SfCHS2-q-F2/R2) flanking the intron21 were designed to evaluate the wild-type transcripts of *SfCHS2* in SS, Sfru_R3, F7-S and F7-R. Parallelly, a pair of primers (SfCHS2-q-F1/R1) that located at the 20th and 21th exon were designed to evaluate the relative expression level of total transcripts of *SfCHS2* in SS, Sfru_R3, F7-S and F7-R.The reaction condition for RT-qPCR was identical with that mentioned above. The relative expression level of *SfCHS2* in Sfru_R3 and individuals from F7-S and F7-R were normalized to the fold value of $2^{-\Delta Ct}$ relative to that in SS.

**CRISPR/Cas9 knockouts**

The CRISPR/Cas9 system was used to create deletions in the *SfCHS2* gene from the SS strain. Briefly, three single-guide RNAs (sgRNA3, sgRNA5 and sgRNA6) were designed using the sgRNAcas9 (V3.0) software (www.biootools.com/software). The template DNA was made with PCR-based fusion of two oligonucleotides with the T7 promoter (Target F: 5'-TAA TAC GAC TCA CTA TAG + the target sequence; Target R: 5'-TTC TAG CTC TAA AAC + the target sequence reverse complement). The target sequences plus PAMs were as follows: sgRNA3 targeting exon 3: sf-chs-sgR3+PAM = 5'-GGA TCT GCG GTT GTG TCT AA GGG-3'; sgRNA5 targeting exon 5: sf-chs-sgR5+PAM = 5'-GTC GTC TGG CCT CTG CTA AA AGG-3', and sgRNA6 targeting exon 6: sf-chs-sgR6+PAM = 5'-AGA CTC GTT ACT ACA CAC AG AGG-3'. For *Spodoptera litura*, the target sequences plus PAMs were sgRNA3 targeting exon 3: Slit_v3-sgR3+PAM = 5'-GGA TCG GCG GTT GTG TCT AA GGG-3' and sgRNA4 targeting exon 4: Slit_v3-sgR+PAM = 5'-AGA GCG TGT GAC ATG GCT GT GGG-3'. For *Mythimna separata*, the target sequences plus PAMs were sgRNA4 targeting exon 4: Msep_v3-sgR4+PAM = 5'-GCG TAT TGG TTT CTC TCG GC GGG-3' and sgRNA5 targeting exon 5: Msep_v3-sgR5+PAM = 5'-CCA TTG CAA AAT CTC CGC GA GGG-3'.*In vitro* transcription was performed with the GeneArt Precision gRNA Synthesis Kit (Thermo Fisher Scientific, Waltham, MA, USA) according to the manufacturer's instructions. The Cas9 protein (GeneArt Platinum Cas9 Nuclease) was purchased from Thermo Fisher Scientific.

For embryo collection and microinjection, freshly laid eggs (within 2 h after oviposition) were immersed in 1% sodium hypochlorite solution for 10 s and washed off from the oviposition gauze, and finally rinsed with distilled water. The eggs were placed on a microscope slide and fixed with double-sided adhesive tape. About 2 nL of a mixture of sgRNAs (250 ng/μL) and Cas9 protein (150 ng/μL) was injected into individual eggs using the Nanoject III (Drummond Scientific, Broomall, PA, USA). The microinjection was completed within 2 h.

Genomic DNA was isolated from a hind leg of $G_0$ adults prior to mating. Primer pairs used to detect specific deletions were as follows: sf-chs-F1 5'-AGC TCA AGA GGC AAA AGG AT-3' in exon 3 and sf-chs-R1 5'-AGC TAA TTG AGT GGC TCC CT-3' in intron 5 for SfCHS2-KO-B; sf-chs-F2 5'-GCC TTC GTA GTA GAC ACC CT-3' in exon 5 and sf-chs-R2 5'-CAT GAA CTT TGT AGA AGC GCT C-3' in exon 6 for SfCHS2-KO-A; and sf-chs-F1 5'-AGC TCA AGA GGC AAA AGG AT-3' in exon 3 and sf-chs-R2 5'-CAT GAA CTT TGT AGA AGC GCT C-3' in exon 6 for SfCHS2-KO-C. For *S. litura*, the primer pairs used to detect deletions for SlCHS2-KO were Slit_v3-F1 5'-AGG ATG GAA TCT GTT CCG AG-3' in exon 3 and Slit_v3-R1 5'-TTG CAA AAA CGT AGG CTT CG-3' in exon 4. Subsequently, PCR products of the region surrounding the target sites were sequenced to determine the exact mutation types in the $G_0$. Selected $G_0$ mutants were crossed to SS to generate $G_1$ progeny. Ultimately, 13 homozygotes with SfCHS2 deletions were mated to establish a homozygous *SfCHS2* knockout strain designated SfCHS2-KO-A.

**Detection of resistance allele of *SfCHS2* with Yaoer insertion in the laboratory and field population of *S. frugiperda***

To identify the Yaoer insertion in the laboratory and field population of *S. frugiperda*, the Whole Genome Sequencing (WGS) reads from 540 samples collected globally were aligned with insertion site sequences. These sequences consist of 600 base pair, representing 300 base pair extensions around the insertion sites. The alignment was performed using BWA. Insertion sites covered by more than three reads were classified as positive insertions. The detailed information for all samples were summarized in (1, 2).

**Hematoxylin and eosin (H&E) staining of midgut tissue section**

The 4th-instar larvae were dissected and the midgut was removed and fixed in mixed picric acid solution. After dehydration, clearing, and paraffin infiltration and paraffin embedding, the samples were cut at 3 μm thickness. Then the sections were deparaffinized, rehydrated, and stained with hematoxylin and eosin (H&E) staining solution (3). The stained sections were scanning using NanoZoomer® S360 microscope (Hamamatsu Photonics Company).

**Subcellular location of transiently expressed SfCHS2 in mammalian and insect cells**

The codon-optimized cDNA of *SfCHS2* was synthesized (GenScript, Nanjing, China) and ligated into the pcDNA3.1(+)-C-DYK vector (GenScript, Nanjing, China) to construct pcDNA-SfCHS2-Flag. The pIE2-EGFP-N1 vector was used to construct plasmid for SfCHS2 overexpression in insect cells. The open reading frame (ORF) of *SfCHS2* was amplified by PCR with cDNA from midgut of SS larvae and ligated into pCE2 TA/Blunt-Zero (Vazyme, Nanjing, China). Then, pIE2-SfCHS2-EGFP was constructed by homologous recombination using ClonExpress® Ultra One Step Cloning Kit (Vazyme, Nanjing, China).

The HEK293T cell line (Human Embryonic Kidney 293 cells) was used for SfCHS2 overexpression. The cells ($5 \times 10^4$ cells/well) were plated onto a cover glass in a 48-well plate (Corning, NY, USA) with complete DMEM (Thermo Fisher Scientific, Waltham, MA, USA) containing 10% heat-inactivated Fetal bovine serum, 100 μg/mL penicillin (Thermo Fisher

Scientific, Waltham, MA, USA), and 100 μg/mL streptomycin (Thermo Fisher Scientific, Waltham, MA, USA). 24 h later, 250 ng pcDNA-SfCHS2-Flag was transfected into cells in each well with 1μL FuGENE Transfection Reagent (Promega, WI, USA) according to the supplier's instructions. 36 h post transfection, the cells were fixed in 4% paraformaldehyde, permeabilized with PBS containing 1% Triton X-100, and blocked with PBS containing 1% BSA. Cells were incubated with a monoclonal antibody against Flag (Sigma, Shanghai, China) diluted in PBS (1:1000) containing 1% bovine serum albumin (BSA) and stained with DyLight 549 secondary antibody (Abbkine, China) at a dilution of 1:1000. Then the cells were stained with 1 μg/mL Hoechst 33258 (Beyotime, Shanghai, China) for 10 min. The images were captured with laser confocal microscope (Leica TCS SP8, Germany). The pre-immune serum was used as a negative control to measure background or cross-reaction. The method is described in details in (4).

*Spodoptera frugiperda* Sf9, or *Trichoplusia ni* Hi5 cells were grown in Grace's insect cell culture medium (Gibco, Thermo, USA) with 10% FBS and 1% penicillin-streptomycin at 28 °C. They were also cultured on a cover glass in 48-well plates (Corning, NY, USA) with complete medium. 24 h later, 250 ng pIE2-SfCHS2-EGFP was transfected into cells in each well with 1 μL FuGENE Transfection Reagent according to the supplier's instructions. 36 h post transfection, the cells were fixed in 4% paraformaldehyde, then stained using 1μg/mL Hoechst 33258 (Beyotime, Shanghai, China) for 10 min. The images were captured with a laser confocal microscope (Leica TCS SP8, Germany).

**Cytotoxicity of activated Vip3Aa toxin to cells**

Three types of cells were cultured in a 48-well plate. 24 h later, HEK293T cells were transfected with pcDNA-SfCHS2-FLAG; Hi5 and Sf9 cells were transfected with pIE2-SfCHS2-GFP. In a control group, the cells were transfected with empty vector. 36 h post transfection, the cells were treated with activated Vip3Aa toxin (diluted in Hanks solution) at various concentrations for 2 h. The images were captured under an inverted microscope (Nikon, Japan).

**Fig. S1. Sample preparation strategy for bulked segregant analysis.** For resistance mapping, a single-pair cross was conducted between a male moth from the Sfru_R3 colony and a female moth from the SS colony to generate F1 progeny. The F1 progeny were raised on a normal diet, resulting in the production of F2 progeny. A total of 960 neonate larvae (480 for both selections) from the F2 generation were subjected to two different diets: a high Vip3Aa diet (4.0 µg/cm$^2$) and a low Vip3Aa diet (0.1 µg/cm$^2$) for a duration of five days. In the case of high Vip3Aa concentration, the individuals that developed into the 3rd instar after 5 days of exposure were classified as resistant to Vip3Aa (F2-R) (n=89). Conversely, for the low Vip3Aa concentration, the individuals that were still <3rd instar were considered susceptible to Vip3Aa (F2-S) (n=75). Following this classification, both the resistant and susceptible larvae from F2 generation were transferred to a normal diet until they reached the fifth instar stage.

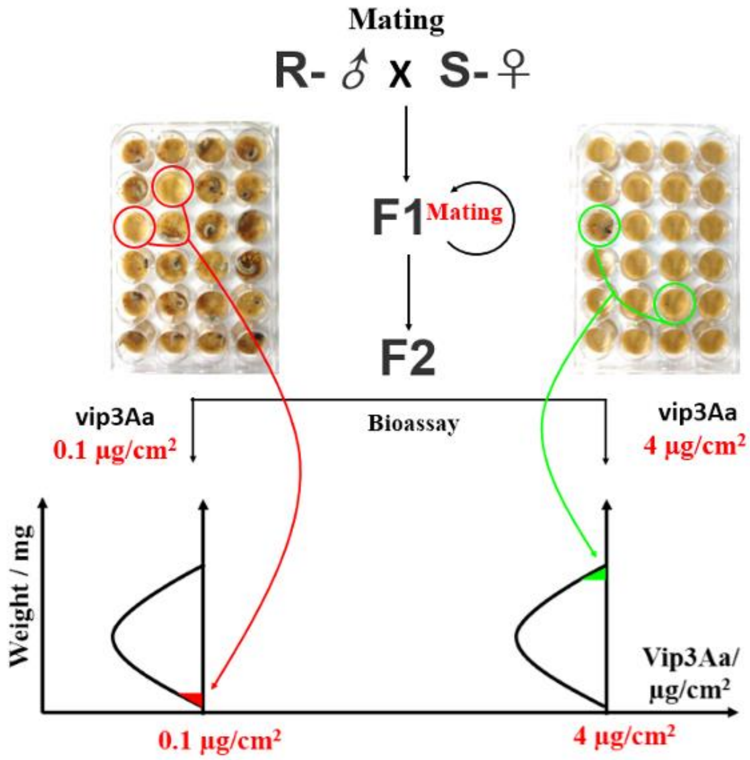

**Fig. S2. Insect chitin synthases.** Protein sequences of chitin synthase 1 (Class A) and chitin synthase 2 (Class B) were aligned using CLUSTAL and a neighbor-joining tree was constructed using MEGAX. Accession numbers are as follows: *Aedes aegypti*, AaCHS1: XP_021704891.1, AaCHS2: XP_001651163.1; *Drosophila melanogaster*, DmCHS1: AAG22215.3, DmCHS2: AAF51798.2; *Manduca sexta*, MsCHS1: AAL38051.2, MsCHS2: AAX20091.1 ; *Spodoptera exigua*, SeCHS1: AAZ03545.1, SeCHS2: ABI96087.1 ; *Spodoptera frugiperda*, SfCHS1: XP_050552783.1, SfCHS2: XP_050552796.1 ; *Tribolium castaneum* TcCHS1: NP_001034491.1, TcCSH2: NP_001034492.

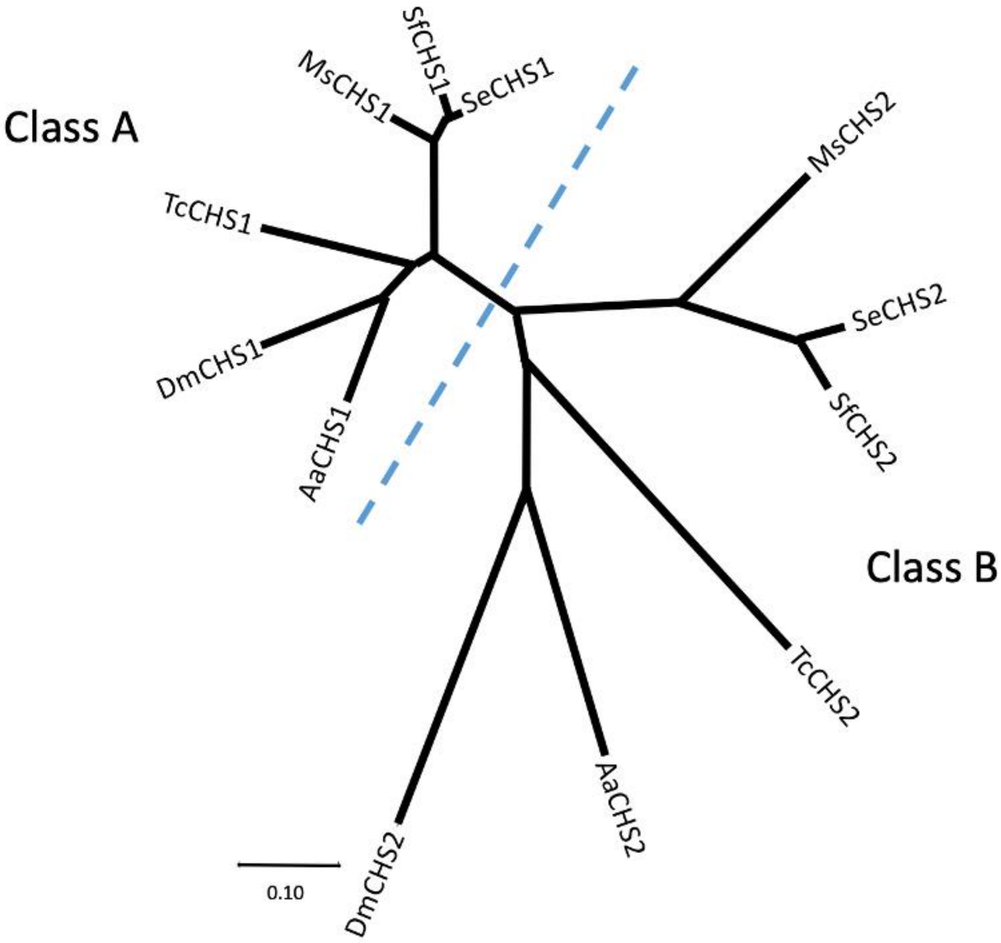

**Fig. S3. Insertion site of LTR retrotransposon Yaoer and alternative splicing of SfCHS2.** (A) By comparison to the wild-type sequence, it is possible to deduce the target site duplication (TSD) typically created by transposable element insertion, which is GAAGG in this case. This represents the last four bases of exon 21 and the first base of intron 21. (B) In the inserted allele, the LTRa immediately follows the first GAAGG, and since the first base of the LTRa is a T, a GT corresponding to the 5'-GU donor site of the pre-mRNA is created immediately after exon 21. The second GAAGG occurs immediately after the second LTR. (C) In accordance with the splicing rules, the entire insert could be spliced out, restoring the wild-type coding sequence (starting with the first G remaining from intron 21, through LTRa, Yaoer, LTRb and to the end of wild-type intron 22 with its 3'-AG acceptor site). (D) However, the inserted allele has the target site duplication GAAGG immediately after the second LTR, immediately preceding the T which is the second base of the wild-type intron 21. This restores the original 5' donor splice site of intron 21. Therefore if just the original intron 21 were spliced out, removing its 3'-AG acceptor site, this could block further splicing of the LTR retrotransposon from the pre-mRNA. In that case, read-through from exon 21 in the mature mRNA would encounter an in-frame stop codon in LTRa, leading to translation of a truncated protein of 1355 residues, where the last 12 residues (VSILFTLLIYLL*) are encoded by the LTR. (E) Alternatively, splicing out just the entire Yaoer element, if possible, would result in another truncated protein ending in VS* due to read-through from exon 21 to intron 21.

(A) pre-insertion state

```
                  GTGAGTTAACTATCTTGA... TTTCCTTTTCGCAG
TGGTATTTCAGTAAGAAG                                   CCGGACGACCTA
 W  Y  F  S  K  K                                     P  D  D  L
   exon 21                    intron 21                 exon 22
```

(B) post-insertion state (TSD = target site duplication)

```
                  GTGTCTATTCTGTTTACATTACTTATTTATTTACTATGACAA...
TGGTATTTCAGTAAGAAG
 W  Y  F  S  K  K
   exon 21         LTRa start

                 TSD
                     GTGAGTTAACTATCTTGA... TTTCCTTTTCGCAG
...TTGGGACGTGAACAGAAG                                   CCGGACGACCTA
                                                         P  D  D  L
      LTRb end                intron 21                    exon 22
```

(C) splicing out the entire insert (GT- from LTRa, -AG from end of intron 21) yields the wild-type protein:

```
TGGTATTTCAGTAAGAAGCCGGACGACCTA ...
 W  Y  F  S  K  K  P  D  D  L
   exon 21          exon 22
```

(D) splicing out just the original intron 21 (GT- from start of intron 21, -AG from end of intron 21) yields a truncated protein:

```
                  GTGTCTATTCTGTTTACATTACTTATTTATTTACTATGACAA...
TGGTATTTCAGTAAGAAG
 W  Y  F  S  K  K  V  S  I  L  F  T  L  L  I  V  L  L  *
    exon 21            LTRa start
```

(E) splicing out just the Yaoer element (GT- from LTRa, -AG from the TSD) also yields a truncated protein:

```
                    GTGAGTTAACTATCTTGA
TGGTATTTCAGTAAGAAG
 W  Y  F  S  K  K  V  S  *

   exon 21          intron 21
```

**Fig. S4. Predicted structures of wild-type and mutant SfCHS2.** The domain structure and the numbers of residues in transmembrane domains and intervening loops were predicted using Phobius (https://phobius.sbc.su.se/). Transmembrane domains are numbered in red. The diagram is not to scale. (A) Full-length wild-type SfCHS2 protein. The C-terminal domain C7 had been used as the bait in a yeast two-hybrid screen for interacting proteins (5). (B) Truncated form (Sfru_R3 allele) produced by insertion of the Yaoer element and read-through from exon 21 to the LTR. Only the C-terminal lumenal domain is affected.

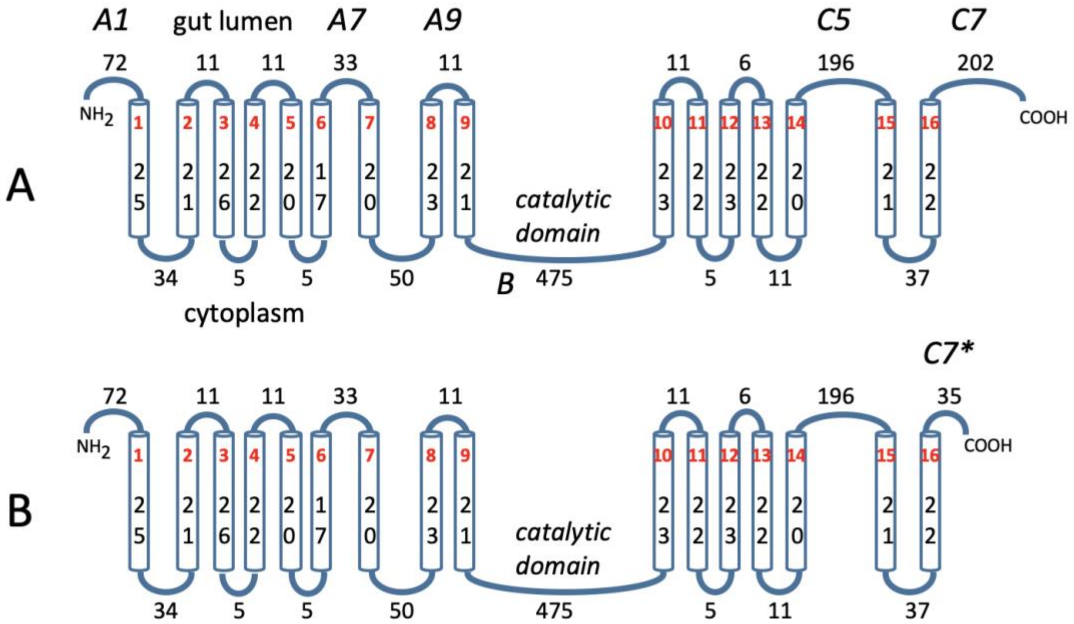

**Fig. S5. Transcripts of *SfCHS2* in SS and Sfru_R3.** (A) *SfCHS2* transcript detection via PCR and Sanger sequencing. Both wild-type and mutant *SfCHS2* transcripts with Yaoer were detected in Sfru_R3. Furthermore, the transcripts including both Yaoer and intron21 were detected in Sfru_R3 by amplification products generated by primer pairs of 11F7/ Nei R5 and Nei F2/11 R4. (B) *SfCHS2* transcripts detection via ISO-seq. In SS, multiple *SfCHS2* isoforms without Yaoer were detected. In contrast, both *SfCHS2* isoforms with and without Yaoer were detected in Sfru_R3. In addition, the isoforms of *SfCHS2* that would have spliced out the Yaoer element while including the intron21 were undetected in Sfru_R3.

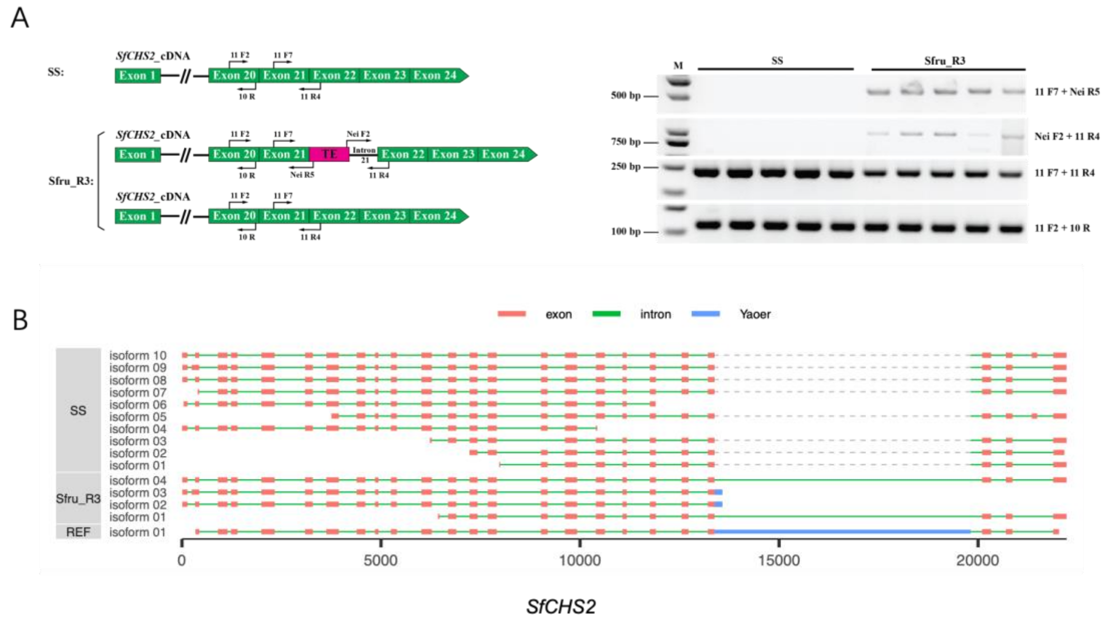

**Fig. S6. Mutation types of *SfCHS2* in SfCHS2-KO-B (A) and SfCHS2-KO-C (B).** CRISPR/Cas9-mediated double sgRNA system and various types of mutations in G1 larvae identified through sequencing of individual PCR clones. Deleted bases are indicated as red dashes, and inserted bases are indicated as red letters. The CRISPR target sites and the number of deleted and inserted bases (+, insertion; –, deletion) are shown. The chromatogram shows the sequence of the mutant isolated from a homozygous knockout larva in G2.

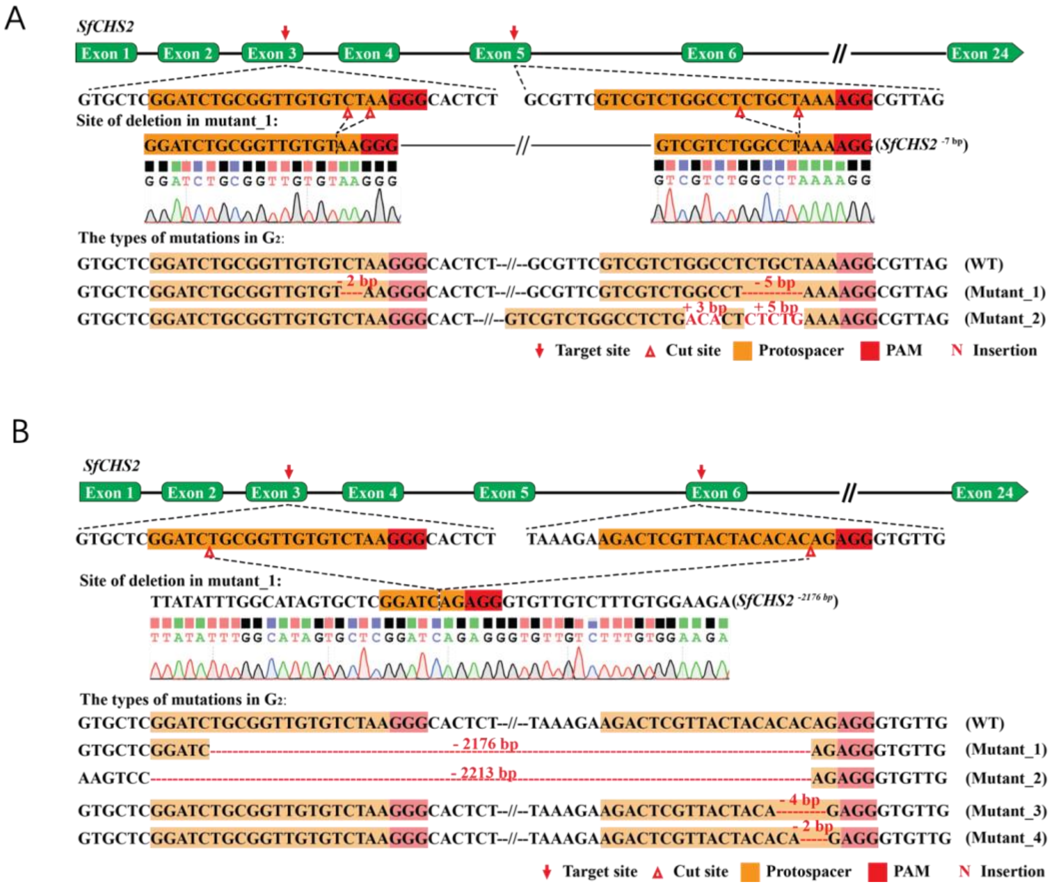

**Fig. S7. Mutation types of *CHS2* in *Spodoptera litura* (A) and *Mythimna separata* (B) via CRISPR/Cas9.** CRISPR/Cas9-mediated double sgRNA system and various types of mutations in G1 larvae identified through sequencing of individual PCR clones. Deleted bases are indicated as red dashes, and inserted bases are indicated as red letters. The CRISPR target sites and the number of deleted and inserted bases (+, insertion; –, deletion) are shown. The chromatogram shows the sequence of the mutant isolated from a homozygous knockout larva in G2.

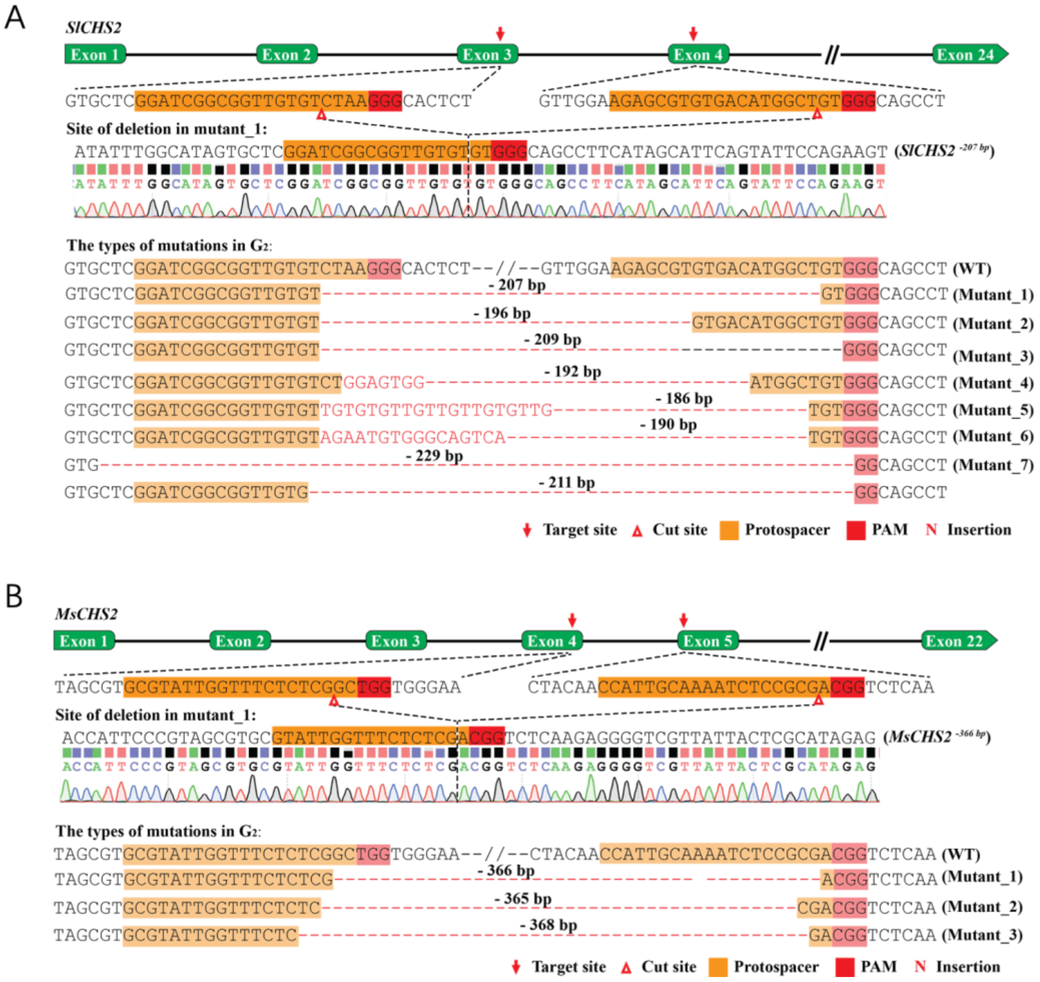

**Fig. S8. Positive mapping of junction of Yaoer and SfCHS2 in an individual collected from the field population of China.** (A) Reads coverage in a sample with a confirmed Yaoer insertion. (B) Reads coverage in a wild-type sample. The insertion site sequences are depicted as colored bars, while the reads are displayed as gray (from R2) and white (from R1) bars. Colored vertical bars within the reads indicate variants.

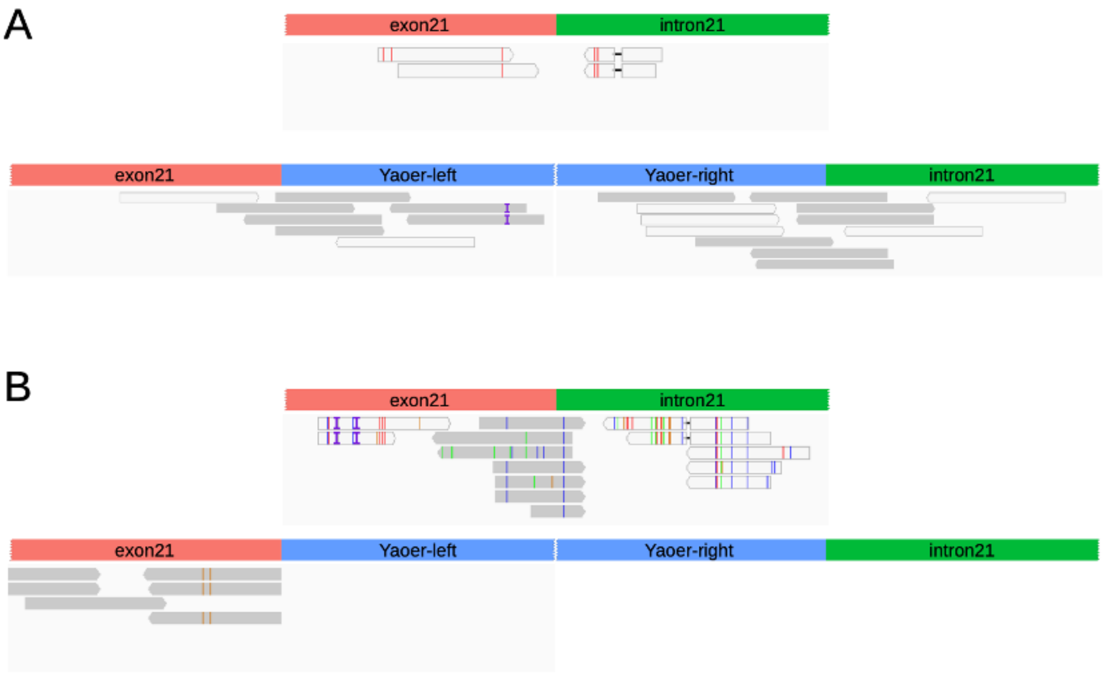

**Fig. S9. Subcellular localization of SfCHS2 in mammalian and insect cells and cytotoxicity test of Vip3Aa toxin.** The pcDNA3.1(+)-C-DYK and pIE2-EGFP-N1 expression vectors were used to construct the recombinant plasmids for heterologous expression of SfCHS2 in HEK 293T, Hi5 and Sf9 cells. (A) The recombinant SfCHS2 were detected mainly in the cytoplasm of all three test cells. (B) No significant phenotypic changes were observed in SfCHS2-overexpressing Hi5 and HEK293T cells after incubation with Vip3Aa activated toxin for 1h. However, the swelling phenotype was observed in non-transfected Hi5 cells when the concentration of Vip3Aa toxin increased up to 80 μg/mL.

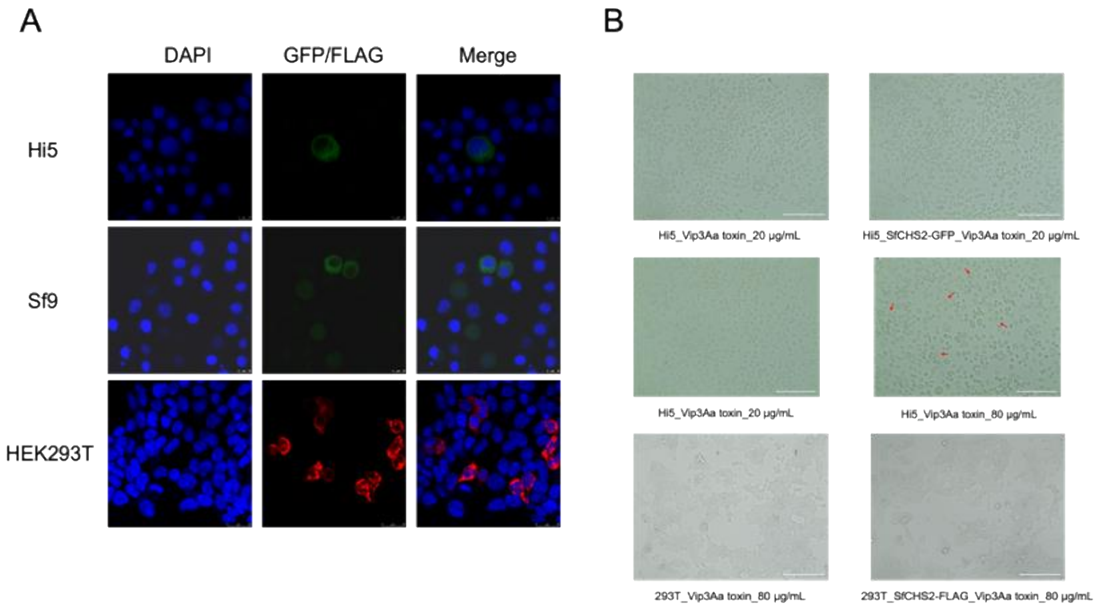

**Table S1. Vip3Aa bioassays during selection process of Sfru_R3 strain.**

| Generation | N[a] | $EC_{50}$ (95% CI) (μg/cm$^2$)[b] | Slope ± SE | $\chi^2$ | df | Resistance ratio [c] | Concentration of Vip3Aa in diet (μg/g) | Date |
|---|---|---|---|---|---|---|---|---|
| 1 | 144 | 0.077 (0.062 - 0.092) | 4.72 ± 0.91 | 0.59 | 3 | - | 2.22 | 2020.06 |
| 3 | 384 | 0.85 (0.30 - 3.67) | 8.39 ± 0.88 | 38.14 | 5 | 11.04 | 2.22 | 2020.08 |
| 12 | 384 | 0.57 (0.33 - 0.98) | 8.21 ± 0.85 | 14.3 | 5 | 7.40 | 4.44 | 2021.05 |
| 13 | 192 | 16.19 (11.22 - 32.70) | 10.57 ± 2.57 | 9.17 | 5 | 210.26 | 8.88 | 2021.07 |
| 14 | 192 | > 16.00 | - | 5.55 | 5 | > 207.79 | 8.88 | 2021.08 |
| 15 | 144 | > 80.00 | - | 2.51 | 3 | > 1038.96 | 12.00 | 2021.09 |
| 16 | 144 | > 100.00 | - | 2.14 | 3 | > 1298.70 | 24.00 | 2021.11 |
| 17 | 64 | 428.26 (312.41 - 616.95.00) | 4.37 ± 1.12 | 0.25 | 2 | 5561.82 | 24.00 | 2021.12 |

[a] Total number of neonates assayed.
[b] Median effective concentration ($EC_{50}$) causing 50% of neonates (first-instar) to die or fail to advance to the third instar in the 7-day test period.
[c] Resistance ratio for an insect population was calculated using its $EC_{50}$ value of the population divided by the $EC_{50}$ of SS.

**Table S2. Survival from first instar to adult eclosion of *S. frugiperda* strains Sfru_R3 and SS on Bt and non-Bt maize.**

| Maize | Strain | Survival (%)[a] | SE (%) | Relative survival [b] |
|---|---|---|---|---|
| Non-Bt | SS | 60.65 | 1.00 | 1.00 |
| | Sfru_R3 | 83.80 | 1.51 | 1.38 |
| Vip3Aa | SS | 0.00 | 0.00 | 0.00 |
| | Sfru_R3 | 23.15 | 4.46 | 0.38 |
| Cry1Ab | SS | 1.39 | 0.65 | 0.02 |
| | Sfru_R3 | 3.70 | 1.36 | 0.06 |
| Cry1Ab + Vip3Aa | SS | 0.00 | 0.00 | 0.00 |
| | Sfru_R3 | 0.00 | 0.00 | 0.00 |

[a] Mean based on three replicates with 72 larvae per replicate (216 larvae per value for survival).
[b] Survival on Bt maize divided by survival on non-Bt maize for the SS strain.

**Table S3. Effective dominance level ($D_{ML}$) of Vip3Aa resistance in *Spodoptera frugiperda* based on the larval mortality observed in the diet-overlay bioassays.**

| Vip3Aa concentration (μg/cm$^2$) | Effective dominance ($D_{ML}$) | Conclusion |
|---|---|---|
| 0.25 | 0.61 | Incomplete dominant |
| 0.5 | 0.45 | Incomplete recessive |
| 1 | 0.065 | Incomplete recessive |

**Table S4. Absence of cross-resistance to Cry toxins.**

| Bt protein | Insect strain | N[a] | $EC_{50}$ (95% CI) (μg/cm$^2$)[b] | Slope ± SE | $\chi^2$ | df | RR [c] |
|---|---|---|---|---|---|---|---|
| Cry1Ab | SS | 144 | 0.16(0.089 - 0.22) | 1.83 ± 0.36 | 2.86 | 3 | -- |
| | Sfru_R3 | 144 | 0.32(0.15 - 0.58) | 0.99 ± 0.28 | 0.32 | 3 | 2.00 |
| Cry1Ac | SS | 144 | 2.95(1.52 - 17.95) | 1.03 ± 0.31 | 1.30 | 2 | -- |
| | Sfru_R3 | 144 | 2.56(1.53 - 8.22) | 1.23 ± 0.32 | 2.22 | 3 | 0.87 |
| Cry1F | SS | 144 | 0.0071(0.0057 - 0.010) | 3.99 ± 0.92 | 2.56 | 2 | -- |
| | Sfru_R3 | 144 | 0.014(0.009 - 0.13) | 0.44 ± 0.84 | 1.50 | 3 | 1.97 |
| Cry2Ab | SS | 144 | 0.094(0.068 - 0.12) | 2.74 ± 0.47 | 2.54 | 3 | -- |
| | Sfru_R3 | 144 | 0.17(0.12 - 0.23) | 1.84 ± 0.36 | 0.28 | 3 | 1.81 |

[a] Total number of neonates assayed.
[b] Median effective concentration ($EC_{50}$) causing 50% of neonates (first-instar) to die or fail to advance to the third instar in the 7-day test period.
[c] RR: resistance ratio for an insect population was calculated using its $EC_{50}$ value of the population divided by the $EC_{50}$ of SS.

**Table S5. Mendelian inheritance of Vip3Aa resistance in Sfru_R3.**

| Insect strain | Pooled backcross | Pooled $F_2$ |
|---|---|---|
| N [a] | 768 | 384 |
| Observed dead (O) | 410 | 299 |
| Expected dead (E) | 384 | 288 |
| $\chi^2$ | 3.52 | 1.68 |
| *P*-value [b] | > 0.05 | > 0.05 |

| Insect strain | Mortality (%) [c] Vip3Aa of 2 μg/$cm^2$ |
|---|---|
| SS | 100.0 ± 0.0 d |
| Sfru_R3 | 0.0 ± 0.0 a |
| F1a (Sfru_R3♂ × SS♀) | 100 ± 0.0 d |
| F1b (Sfru_R3♀ × SS♂) | 100 ± 0.0 d |
| Pooled F1 | 100.0 ± 0.0 |
| F2a (F1a × F1a) | 76.0 ± 2.9 c |
| F2b (F1b × F1b) | 79.7 ± 3.3 c |
| Pooled F2c | 77.9 ± 2.2 |
| BCR1 (Sfru_R3♂ × F1a♀) | 55.2 ± 2.3 b |
| BCR2 (Sfru_R3♀ × F1a♂) | 52.6 ± 4.4 b |
| BCR3 (Sfru_R3♂ × F1b♀) | 50.0 ± 5.7 b |
| BCR4 (Sfru_R3♀ × F1b♂) | 55.7 ± 2.8 b |
| Pooled backcross [d] | 53.4 ± 2.0 |
| F-test F-value | $F_{9, 76} = 96.79$ |
| *P*-value | < 0.0001 |

[a] Total number of neonates tested.
[b] $\chi^2_{0.05}$（df = 1）= 3.84.
[c] Mean values within a column followed by the same letter are not significantly different at α = 0.05 (Tukey's HSD test). Larval mortality was corrected based on the mortality observed on the control diet.
[d] The pooled data were not included in ANOVA and HSD tests.

**Table S6. Normalized midgut expression levels of genes in the 0.45 Mb interval on Chromosome 1.** FPKM = Fragments Per Kilobase of transcript per Million mapped reads. The GenBank annotations for the chitin synthases are incorrect. (**) = SfCHS2 (class B) chitin synthase 2, (*) = SfCHS1 (class A) chitin synthase 1. Gene ID and GenBank Annotations are from the recent version of the genome of *Spodoptera frugiperda*, AGI-APGP_CSIRO_Sfru_2.0 (GCF_023101765.2).

| Gene ID | Start (Mb) | End (Mb) | FPKM[a] | GenBank Annotation |
|---|---|---|---|---|
| **LOC118273136** | **8,175,293** | **8,181,479** | **3.43** | **geranylgeranyl transferase type-2 subunit alpha-like** |
| LOC118273137 | 8,182,314 | 8,184,055 | 17.15 | protein abrupt-like |
| LOC118273138 | 8,186,422 | 8,467,817 | 0.28 | neural-cadherin-like |
| LOC118273139 | 8,282,422 | 8,284,761 | 0.00 | uncharacterized LOC118273139 |
| LOC118273194 | 8,502,522 | 8,510,565 | 1.02 | formin-like protein 5 |
| LOC118273449 | 8,511,257 | 8,512,953 | 29.36 | probable 26S proteasome non-ATPase regulatory subunit 3 |
| LOC118273513 | 8,513,304 | 8,514,967 | 0.02 | dynein assembly factor 6, axonemal-like |
| LOC118273659 | 8,519,645 | 8,531,980 | 0.28 | cysteine-rich secretory protein 1-like |
| LOC118273515 | 8,534,307 | 8,549,529 | 0.06 | protein phosphatase PHLPP-like protein |
| LOC118273516 | 8,549,313 | 8,589,130 | 0.00 | LIM/homeobox protein Lhx3-like |
| LOC118273715 | 8,589,783 | 8,595,627 | 0.00 | cilia- and flagella-associated protein 47-like |
| LOC118273406 | 8,598,343 | 8,604,305 | 0.00 | 3,4-dihydroxyphenylacetaldehyde synthase 2-like |
| LOC118273430 | 8,604,300 | 8,605,505 | 1.60 | uncharacterized protein LOC118273407 |
| LOC118273345 | 8,609,025 | 8,610,950 | 9.69 | negative elongation factor E-like |
| LOC118273453 | 8,611,248 | 8,614,886 | 8.87 | cysteine desulfurase, mitochondrial-like |
| LOC118273454 | 8,614,789 | 8,616,366 | 6.54 | ras-related protein Rab-1A-like |
| LOC118273568 | 8,618,802 | 8,623,616 | 0.48 | MPN domain-containing protein CG4751-like |
| **LOC118273105** | **8,624,065** | **8,639,856** | **111.97** | **chitin synthase chs-2-like (**)** |
| LOC118273150 | 8,641,476 | 8,668,929 | 1.25 | chitin synthase chs-2-like (*) |

[a] The FPKM value refers to Fragments Per Kilobase of transcript per Million mapped reads.

**Table S7. Susceptibility of *SfCHS2*-knockout strains and progeny of crosses with Sfru_R3 to Vip3Aa toxin.**

| Insect strain | N[a] | $EC_{50}$ (95% CI) (μg/cm$^2$)[b] | Slope ± SE | $\chi^2$ | df | RR [c] |
|---|---|---|---|---|---|---|
| SS | 192 | 0.13 (0.11 - 0.16) | 3.98 ± 0.62 | 6.39 | 5 | - |
| SfCHS2-KO-A | 80 | > 1600 | - | - | - | > 12307.69 |
| SfCHS2-KO-B | 96 | > 1600 | - | - | - | > 12307.69 |
| SfCHS2-KO-C | 96 | > 1600 | - | - | - | > 12307.69 |
| SfCHS2-KO♀ × Sfru_R3♂ | 80 | > 1600 | - | - | - | > 12307.69 |
| SfCHS2-KO♂ × Sfru_R3♀ | 80 | > 1600 | - | - | - | > 12307.69 |

[a] Total number of neonates assayed.

[b] Median effective concentration ($EC_{50}$) causing 50% of neonates (first-instar) to die or fail to advance to the third instar in the 7-day test period.

[c] Resistance ratio for an insect population was calculated using its $EC_{50}$ value of the population divided by the $EC_{50}$ of SS.

**Table S8. Susceptibility of *CHS2*-knockout strains of *S. litura* and *M. separata* to Vip3Aa toxin.**

| Insect strain | N[a] | $EC_{50}$ (95% CI) ($\mu g/cm^2$) [b] | Slope ± SE | $\chi^2$ | df | Resistance ratio [c] |
|---|---|---|---|---|---|---|
| Sl-SS | 96 | 0.016 (0.0030 - 0.086) | 4.08 ± 0.82 | 14.45 | 5 | -- |
| SlCHS2-KO | 96 | >1600 | 0 | -- | -- | > 100,000 |
| Ms-SS | 168 | 1.20 (0.97 - 1.48) | 6.56 ± 1.08 | 2.93 | 4 | -- |
| MsCHS2-KO | 96 | >1600 | 0 | -- | -- | > 1,333.3 |

[a] Total number of neonates assayed.

[b] Median effective concentration (EC50) causing 50% of neonates (first-instar) to die or fail to advance to the third instar in the 7-day test period.

[c] Resistance ratio for an insect population was calculated using its $EC_{50}$ value of the population divided by the $EC_{50}$ of SS.

**Table S9. Fitness parameters and survival on artificial diet.** Larvae in Vip3Aa treatments were fed on artificial diet treated with Vip3Aa (24 µg/g) until pupation.

| **Artificial Diet** | **Larval stage duration in days (Mean ± SE)** | |
|---|---|---|
| Strain | female | male |
| SS | 13.39 ± 0.093 | 13.48 ± 0.11 |
| Sfru_R3 | 14.10 ± 0.050 | 13.98 ± 0.066 |
| Sfru_R3 on Vip3Aa | 16.39 ± 0.12 | 16.45 ± 0.079 |
| SfCHS2-KO-A | 13.37 ± 0.088 | 13.37 ± 0.094 |
| SfCHS2-KO-A on Vip3Aa | 19.49 ± 0.20 | 20.26 ± 0.26 |
| **Artificial Diet** | **Pupal stage duration in days (Mean ± SE)** | |
| Strain | female | male |
| SS | 8.74 ± 0.070 | 10.09 ± 0.048 |
| Sfru_R3 | 10.11 ± 0.074 | 11.08 ± 0.056 |
| Sfru_R3 on Vip3Aa | 10.13 ± 0.088 | 11.65 ± 0.076 |
| SfCHS2-KO-A | 9.947 ± 0.037 | 11.34 ± 0.075 |
| SfCHS2-KO-A on Vip3Aa | 9.92 ± 0.069 | 10.93 ± 0.054 |
| **Artificial Diet** | **Pupal weight in mg (Mean ± SE)** | |
| Strain | female | male |
| SS | 203.09 ± 3.94 | 217.07 ± 3.61 |
| Sfru_R3 | 212.42 ± 3.74 | 225.88 ± 3.91 |
| Sfru_R3 on Vip3Aa | 202.35 ± 3.97 | 205.36 ± 2.44 |
| SfCHS2-KO-A | 233.47 ± 3.83 | 239.95 ± 2.80 |
| SfCHS2-KO-A on Vip3Aa | 135.51 ± 3.06 | 148.03 ± 3.67 |

**Table S10. Partial primers used in this study.**

| Primer name | Oligonucleotide (5'-3') | Application |
|---|---|---|
| 2992-SNP-F | AGTGCTAAAGAGGGCTAAGG | Fine-scale mapping |
| 2992-SNP-R | TCCAGGTAATCCGACACTGGG | |
| 3106-SNP-F | AGGTCAGTGTCCCACTAC | |
| 3106-SNP-R | ATGGCACTTCTACCGTTGG | |
| 3136-SNP-F1 | ACTATAGATAGCTGGAGAGCC | |
| 3136-SNP-R1 | ACGAAGCTAACACATTAGACG | |
| 3105-SNP-F2 | AGCTCTTCCTCGCTAACCTC | |
| 3105-SNP-R2 | TTTCTTCTGCGGCGTCTCTC | |
| 3493-SNP-F | TGTTCGATGGAGGACATGGTC | |
| 3493-SNP-R | TGTAAATGAAACCGCCAGCAG | |
| 3409-SNP-F1 | AGATGGGAAGATGAGAGACCC | |
| 3409-SNP-R1 | ACCTCTCCCTGATGAAAGAAG | |
| 3178-SNP-F1 | TGTACCTGAAGTACCAGCTCC | |
| 3178-SNP-R1 | CTGTGGCTAGGTTCTACTTCC | |
| 3547-SNP-F1 | ATTGGAGCGCAACGTTGAGTTG | |
| 3547-SNP-R1 | AGGTGTTCCAGTTGTGGATG | |
| 11F7 | CGGTTTCGTGTTCCTCCTGT | Transcripts detection |
| NeiR5 | GACCGCTGTGTACCACTTGT | |
| NeiF2 | TCGGGTGGTAACAATACGGT | |
| 11R4 | AGTCGTCGGTGTTCAGACGT | |
| 11 R4 | AGTCGTCGGTGTTCAGACGT | |
| 11 F2 | GGTGTTTGCGTTCGTGATGT | |
| 10 R | GTCGTACTCCATGCTGGACT | |

**SI References**

1. L. Zhang *et al.*, Global genomic signature reveals the evolution of fall armyworm in the Eastern hemisphere. *Mol. Ecol.* **32**, 5463-5478 (2023).
2. L. Zhang *et al.*, Genetic structure and insecticide resistance characteristics of fall armyworm populations invading China. *Mol. Ecol. Resour.* **20**, 1682-1696 (2020).
3. A. T. Feldman, D. Wolfe, Tissue processing and hematoxylin and eosin staining. *Methods Mol. Biol.* **1180**, 31-43 (2014).
4. Z. Gai *et al.*, Characterization of Atg8 in lepidopteran insect cells. *Arch. Insect Biochem. Physiol.* **84**, 57-77 (2013).
5. G. Broehan, L. Zimoch, A. Wessels, B. Ertas, H. Merzendorfer, A chymotrypsin-like serine protease interacts with the chitin synthase from the midgut of the tobacco hornworm. *J. Exp. Biol.* **210**, 3636-3643 (2007).
